# Dual MYC and GSPT1 Protein Degrader for MYC-Driven Cancers

**DOI:** 10.1101/2025.04.24.650490

**Authors:** Yuki Nishida, Valeria Impedovo, Edward Ayoub, Natalia Baran, Darah A. Scruggs, Hideaki Mizuno, Shayaun Khazaei, Lauren B. Ostermann, Sandeep Singh, Andrea Bedoy, Po Yee Mak, Bing Z. Carter, Eiji Sugihara, Tetsuya Takimoto, Youzhi Tong, Qianxiang Zhou, Zhaohui Yang, Honghua Yan, Dong Chen, Abhishek Maiti, Koji Sasaki, Steffen Boettcher, Torsten Haferlach, Stefano Tiziani, Liandong Ma, Michael Andreeff

## Abstract

Direct targeting of the oncoprotein MYC has long been attempted in cancer therapy, with limited success. We here identify a novel co-regulatory feedback loop of MYC and G1 to S phase transition protein 1 (GSPT1), where MYC promotes transcription of GSPT1, and GSPT1 senses stop codon of *MYC* to promote its translation. We report on the first-in-class dual MYC/GSPT1 protein degrader, GT19630. GT19630 significantly induced integrated stress response, abrogated oxidative phosphorylation through inhibition of the TCA cycle and induced cell death. Protein degradation of MYC was critical for efficacy of GT19630. GT19630 induced profound anti-proliferative effects and apoptosis agnostic to *TP53* in a broad range of cancer cells, and is highly active in vivo in multiple, therapy-resistant hematologic and solid tumor models. Dual MYC/GSPT1 degradation was well tolerated in humanized *Crbn^I391V^* mice. In conclusion, we propose a novel treatment approach by directly targeting the MYC-GSPT1 axis in MYC-driven cancers.

**Statement of significance:** MYC has been considered an undruggable protein. We found a targetable, novel positive co-regulatory feedback of MYC and GSPT1, a key translation terminator. The dual MYC/GSPT1 degrader GT19630 is highly active in MYC-driven tumors, with moderate effects on humanized Crbn mice, providing opportunities to improve treatment outcome of MYC-driven cancers.

## Introduction

The oncoprotein c-MYC (hereafter, MYC) governs the transcription of approximately 15% of the human genome and is dysregulated in 70% of all human cancers^1–3^. The translocation of *IgH/MYC* in Burkitt’s lymphoma was the first reported manifestation of MYC driving human cancer^4,5^, followed by identification of *MYC* gene amplification, constitutive activation, overexpression and increased protein stability in other tumors^6,7^. Genetic studies have shown that the ablation of *MYC* exhibits profound anti-proliferative effects with sustained tumor regression in multiple cancer models^8^. MYC controls the maintenance of cancer and leukemia stem cells, conferring resistance to chemo- and targeted therapies^3^. Recent analyses have revealed MYC overexpression in acute myeloid leukemia (AML) stem cell populations^9^. We previously reported that in primary AML samples, MYC overexpression conferred resistance to inhibition of BCL-2, an antiapoptotic protein, and that MYC-induced resistance could be targeted indirectly by reactivation of the tumor suppressor p53^10^. These findings strongly suggest MYC as bona fide target in multiple cancers.

Because of its subcellular localization and intrinsically disordered structure, MYC has long been considered “undruggable,”^2^ and direct targeting of MYC protein has not been reported. Targeted protein degradation is an emerging treatment modality encompassing two approaches, proteolysis targeting chimeras (PROTACs) and cereblon E3 ligase modulators (CELMoDs) ^11,12^. Recent studies have revealed neosubstrates, novel pathogenic molecular targets of protein degraders, especially in myeloid and lymphoid malignancies^13^. One such neosubstrate is the G1 to S phase transition protein 1 (GSPT1) (also known as eukaryotic release factor 3a, eRF3a), a translation termination factor that cooperates with eRF1 encoded by *ETF1*^14^. GSPT1 has been investigated as a molecular target, with promising preliminary efficacy and selectivity associated with its degradation^15–18^. Although the blockade of translation termination reduces protein stability and induces integrated stress response as mode of action for inducing lethality in cancer cells^16,17^, the role of GSPT1 in the stability of MYC protein has not been investigated.

MYC regulates virtually all the essential genes involved in glycolysis and glutaminolysis^19^. Other metabolic processes regulated by MYC include fatty acid and essential amino acid metabolism and nucleotide biosynthesis^19^. The inhibition of MYC, therefore, rewires a multitude of metabolic processes involving glucose, glutamine, essential amino acids and fatty acids ^19,20^. On the other hand, the effects of MYC ablation in different cell types under different conditions are context dependent. For instance, MYC knockdown reduces glycolysis in non-cancerous cells^21^ but was reported to reduce oxidative phosphorylation (OXPHOS) in breast cancer stem cells^22^. The direct metabolic consequences of MYC and GSPT1 protein degradation remain unexplored.

Here ,we report a novel, co-regulatory feedback loop of MYC and GSPT1 involved in transcriptional and translational modulations, and describe the development of a first-in-class dual MYC/GSPT1 degrader that targets this co-regulatory feedback loop. Dual MYC/GSPT1 degradation exhibited profound anti-cancer activities in multiple models of MYC-driven hematologic and solid tumors in vitro and in vivo, with a therapeutic differential between normal hematopoietic stem/progenitor cells (HSPCs) and leukemia stem/progenitor cells (LSPCs) and favorable toxicity profiles in humanized *Crbn*-mutant mice. We discovered that dual MYC/GSPT1 degradation primarily induces profound OXPHOS inhibition and impairs the tricarboxylic acid (TCA) cycle at the glutamate and citrate synthesis levels. These findings pave the way for a novel treatment strategy that directly degrades MYC/GSPT1 protein in MYC-driven cancers.

## Results

### Interdependent roles of MYC and GSPT1 and development of dual MYC and GSPT1 degrader

We first investigated whether MYC regulates translation termination factors *GSPT1* and *ETF1* which encode eRF3a and eRF1, respectively. MYC binds to the transcription start sites of both *GSPT1* and *ETF1* in hematologic and solid tumor cells (**Fig. 1 A**, **Supplementary Fig. S1**). To investigate the interactions of MYC and GSPT1, we first knocked down *MYC* and/or *GSPT1* using siRNAs in MYC-driven AML HL-60 cells harboring the double-minute chromosome abnormality, which results in MYC amplification^23^. *MYC* knockdown reduced GSPT1 mRNA and protein levels (**Supplementary Fig. S2A**), and *GSPT1* knockdown reduced MYC protein levels (**Fig. 1A, Supplementary Fig. S1B**). The double knockdown of *MYC* and *GSPT1* further decreased levels of both proteins (**Fig. 1A**), suggesting that MYC is an upstream regulator of GSPT1, and MYC and GSPT1 form a positive feedback loop. *MYC* and *GSPT1* are amongst the essential genes, particularly in blood cancer cell lines with negative dependency scores upon CRISPR knockout based on DepMap data (**Supplementary Fig. S1C**) ^24^, providing rationale for the hypothesis that dual targeting of MYC and GSPT1 constitutes a highly potent, novel treatment approach, especially for MYC-driven hematologic malignancies.

**Fig. 1.**
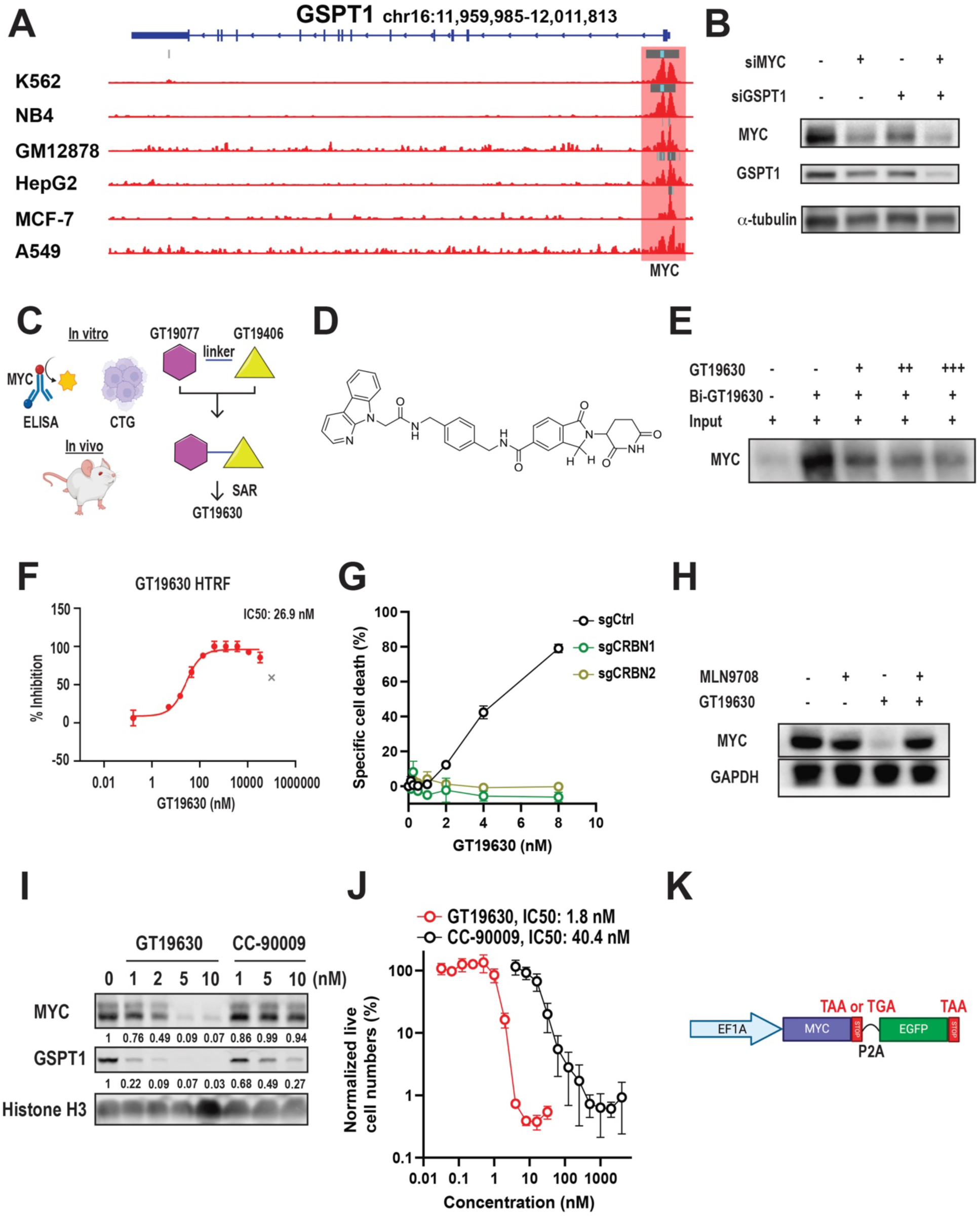
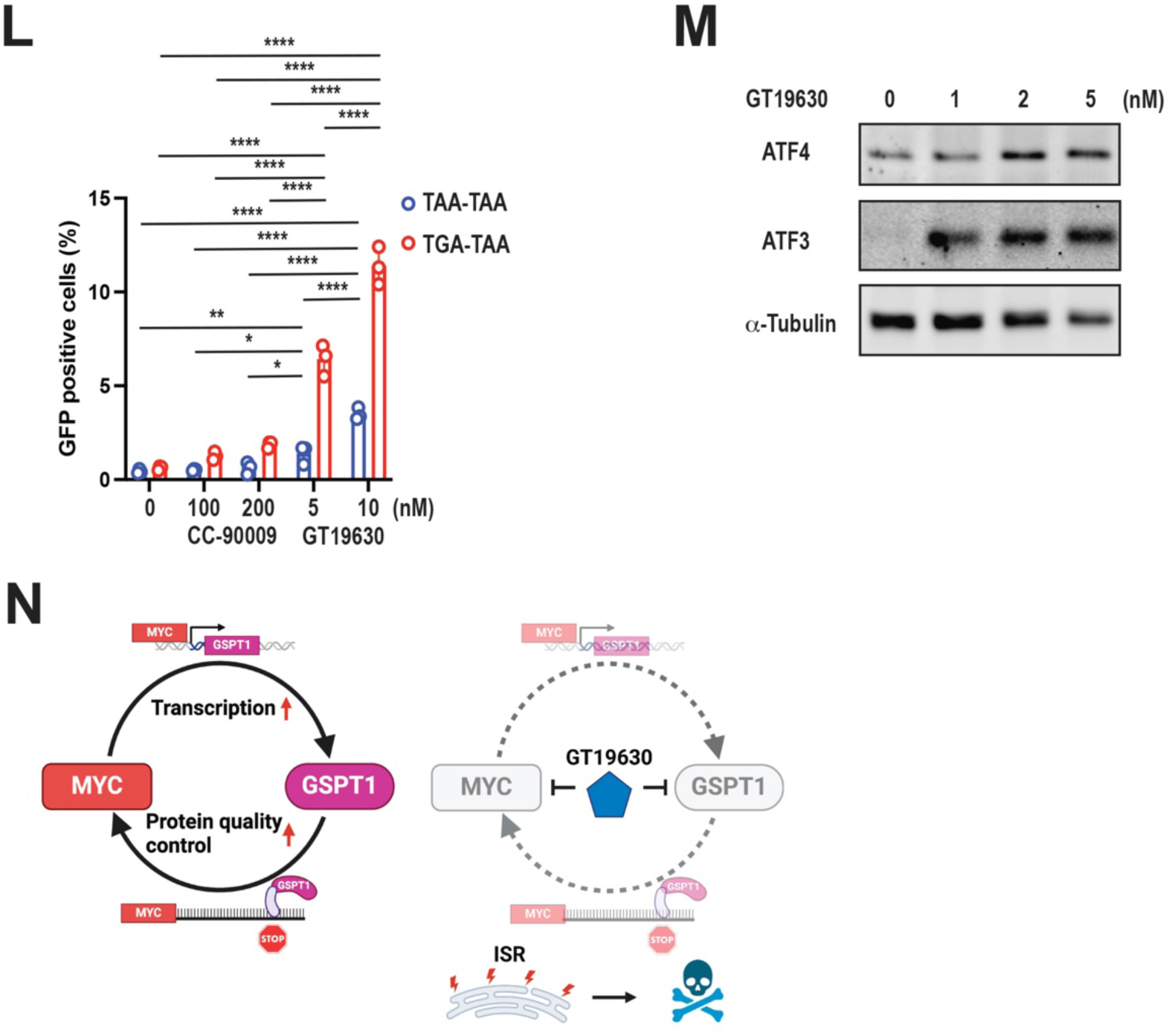
Co-regulatory Feedback Loop of MYC and GSPT1, and discovery of MYC/GSPT1 Degrader. **A.** Binding of MYC to the TSS of GSPT1 in hematologic and solid tumor cells. GSPT1 gene map at chromosome 16 is shown on top. Significantly enriched regions are highlighted with gray bars above peaks. Highest peaks are highlighted in light blue. **B.** Western blot of MYC and GSPT1 in HL-60 cells transduced with small interfering RNA for MYC (siMYC), GSPT1 (siGSPT1) or both siMYC and siGSPT1. α-Tubulin served as loading control. **C.** Schematic diagram of designing and developing MYC/GSPT1 degraders GT19630 and GT19630. **D.** Structure of GT19630. **E.** Western blot after cell-free, affinity pull-down assay for recombinant human MYC protein incubated with biotinylated (Bi)-GT19630 or with increasing doses of naked GT19630. **F.** Homogeneous time-resolved fluorescence (HTRF) assays for the interaction of CRBN with GT19630. **G.** Percentages of specific cell death in Cas9-expressing MOLM-13 cells transduced with the control gRNA (sgCtrl), gRNAs for CRBN (sgCRBN1 and sgCRBN2). **H.** Western blot of MYC after treatment with DMSO, 50 nM MLN9708, 10 nM GT19630 and both MLN9708 and GT19630 in HL-60 cells for 8 hours. GAPDH served as loading control. **I.** Western blot of MYC and GSPT1 in HL-60 cells treated with indicated concentrations of GT19630 and CC-90009 for 24 hours. Histone H3 served as loading control. **J.** Normalized live cell numbers of HL-60 cells treated with indicated concentrations of GT19630 or CC-90009 for 48 hours. **K.** Construct scheme of the MYC-stop-EGFP fluorescent translation termination reporter. The *MYC* gene is followed by two different stop codons (either TAA or TGA), indicated by red rectangles. P2A was used as a linker for the EGFP sequence. **L.** Percentages of GFP-positive HL-60 cells with two different MYC-stop-GFP constructs (TAA or TGA) treated with indicated concentrations of CC-90009 or GT19630 for 24 hours. **M.** Western blot of ATF4 and ATF3 in HL-60 cells treated with indicated concentrations of GT19630 for 24 hours. α-Tubulin served as loading control. **N.** Schematic diagrams of a co-regulatory feedback loop between MYC and GSPT1 (left) and GT19630’s interference (right). Degradation of MYC and GSPT1 leads to integrated stress response (ISR). \**P* < 0.05, \*\**P* < 0.01, \*\*\*\**P* < 0.0001. Representative results of three biological replicates are shown for WB data.

As GSPT1 is a known neosubstrate of cereblon (CRBN)-mediated protein degradation, we aimed to screen for a PROTAC active against MYC and GSPT1. We started targeting MYC by pharmacological inhibition of MYC:MAX protein-protein interaction and developed GT19077 as an inhibitor of their interaction. We then used GT19077 as “warhead” and GT19406 as E3 ligand for CRBN. MYC ELISAs were used as primary cell-based assays to direct structure-activity relationship efforts, resulting in the identification of GT19630 as lead compounds for MYC degradation (**Fig. 1C, D**; further detail is described in **Supplementary Fig. S2A** and the invention disclosure (WO2022268066A1)). Confirming the binding of GT19630 to MYC protein, a large amount of pulled-down MYC protein was observed when incubated with biotinylated GT19630, and the addition of naked (nonbiotinylated) GT19630 reduced the amount of pulled-down MYC protein in a concentration-dependent manner in cell-free affinity pull-down assays (**Fig. 1D**). A homogenous time-resolved fluorescence assay for CRBN binding revealed an approximately 200-fold higher binding affinity of GT19630 with CRBN (IC50 value of 26.9 nM) compared to thalidomide, a classic CELMoD (IC50 value of 5.4 μM) (**Fig. 1E, Supplementary Fig. S2B**), suggesting a promoted CRBN binding affinity. CRISPR knockout of CRBN completely abrogated GT19630-induced cell death in MOLM-13 cells (**Fig. 1F**, **Supplementary Fig. S2C**). GT19630 profoundly reduced MYC protein in HL-60 cells, and the reduction was completely rescued by adding proteasome inhibitors MLN9708 (ixazomib; **Fig. 1G**) or bortezomib (**Supplementary Fig. S2D**), suggesting proteasome-dependent protein degradation of MYC. GT19630 reduced MYC protein levels in a dose- and time-dependent manner (**Supplementary Fig. S2E, F**) at low nanomolar concentrations, and the reduction of MYC was sustained up to 48 hours after washout of GT19630 in HL-60 cells (**Supplementary Fig. S2G**), suggesting high potency and intracellular stability. Interestingly, we found that GT19630 degraded both MYC and GSPT1, while CC-90009, a selective CELMoD for GSPT1, did not degrade MYC (**Fig. 1H**), suggesting that GT19630 possesses characteristics of a molecular glue. GT19630 increased cell death compared to CC-90009, with approximately 20-fold lower IC50 values (**Fig. 1H**). The superior efficacy of GT19630 over CC-90009 was also observed in other AML cell lines (**Supplementary Fig. S2H, I**), suggesting an advantage of both MYC and GSPT1 degradation over GSPT1 degradation alone. These data suggest that GT19630 and GT19630 induce dual degradation of MYC and GSPT1 through binding to both MYC and GSPT1 utilizing a CRBN-mediated proteasome system.

GSPT1 degradation leads to an increased frequency of stop-codon readthrough (SCR) ^16,25^, resulting in increased unfolded peptides and the induction of integrated stress response (ISR) ^16,17^. However, whether GSPT1 controls MYC translation, and more specifically, whether SCR occurs on *MYC* upon GSPT1 reduction, remained unknown. To address this, we established a reporter assay for exogenous *MYC* with either TAA or TGA stop codons followed by green fluorescent protein (GFP) serving as readout of SCR after *MYC* mRNA translation (**Fig. 1J**). As positive control, we used G418, an aminoglycoside antibiotic that induces SCR ^26^. As expected, G418 induced GFP-positive cells, especially in MYC-TGA HL-60 and HEK293T cells, proving the system functioned properly (**Supplementary Fig. S3A-C**). CC-90009 induced only minimal GFP signals, while GT19630 significantly and profoundly induced GFP signals in both HL-60 (**Fig. 1K**) and HEK293T cells (**Supplementary Fig. S3D**) with the TGA stop codon, suggesting that dual MYC/GSPT1 degradation enhances GSPT1 degradation–mediated SCR of *MYC* mRNAs.

Next, we asked whether GT19630 induces integrated stress response upon increased SCR. We generated GT19630-sensitive (GTS) and -resistant (GTR) HL-60 cells through chronic exposure to GT19630 (**Supplementary Fig. S3E**). HL-60 GTR cells expressed lower CRBN levels than HL-60 GTS cells (**Supplementary Fig. S3F**), partially explaining resistance to CRBN-utilizing drugs as shown in multiple myelomas^27^. RNA-sequencing (RNA-seq) and pathway analyses revealed that GT19630 significantly upregulated integrated stress response (ISR), stress response and endoplasmic reticulum stress pathways in three different gene set enrichment analyses of HL-60 GTS cells (**Supplementary Fig. S3G**). GT19630 induced ATF4, the master transcription factor for ISR and its key downstream target ATF3 (**Fig. 1L**).

Taken together, we identified a positive, co-regulatory feedback loop employing transcriptional and translational controls between MYC and GSPT1 and developed the novel, first-in-class dual MYC/GSPT1 degrader GT19630, which induces ISR as the principal mechanism of action (**Fig. 1M**).

### Dual MYC/GSPT1 degradation induces profound OXPHOS inhibition

As MYC governs a wide range of metabolomic pathways in cancer, we interrogated the impact of GT19630 on metabolic changes in HL-60 GTS and GTR cells. First, we performed RNA-seq analyses of HL-60 GTS cells treated with DMSO or GT19630. Pathway analyses of differential gene expression profiles revealed, as expected, that GT19630 significantly downregulated MYC targets. GT19630 also significantly downregulated oxidative phosphorylation (OXPHOS) and fatty acid metabolism, but not glycolysis (**Fig. 2A, Supplementary Fig. S4A**). Supporting this finding, Seahorse analyses of HL-60 GTS cells treated with DMSO, GT19630 or CC-90009 showed significant decreases in basal and maximal oxygen consumption rates (OCRs) induced by GT19630 compared to DMSO or CC-90009 (**Fig. 2B-D**). GT19630 also significantly decreased OCR-linked ATP production and the ratio of OCR and extracellular acidification rate (ECAR) compared to DMSO or CC-90009 (**Fig. 2E, F**), demonstrating the superiority of targeting both MYC and GSPT1 compared to GSPT1 alone. GT19630 also significantly reduced ECAR compared to DMSO or CC-90009 (**Fig. 2G**). These changes observed in HL-60 GTS cells were not observed in HL-60 GTR cells (**Supplementary Fig. S4B-G**). To gain deeper insights into the dynamic metabolomic impact of GT19630, we performed untargeted metabolomic analyses in HL-60 GTS cells treated with DMSO or GT19630. GT19630 significantly reduced levels of glutamine, leading to reduced levels of succinate, malate, alanine and aspartate and increased levels of citrate compared to DMSO control (**Fig. 2H, I**), suggesting that GT19630 inhibited the TCA cycle by disrupting glutamine metabolism. On the other hand, GT19630 significantly increased levels of glucose-6-phosphate (G6P), fructose-1,6-bisphosphate and phosphoenolpyruvate (PEP), while pyruvate levels were not changed (**Fig. 2J, K**), suggesting the accumulation of glycolysis intermediates after inhibition of the TCA cycle. In summary, our findings show that MYC/GSPT1 degradation primarily inhibits OXPHOS and mitochondrial respiration through disruption of the TCA cycle.

**Fig. 2.**
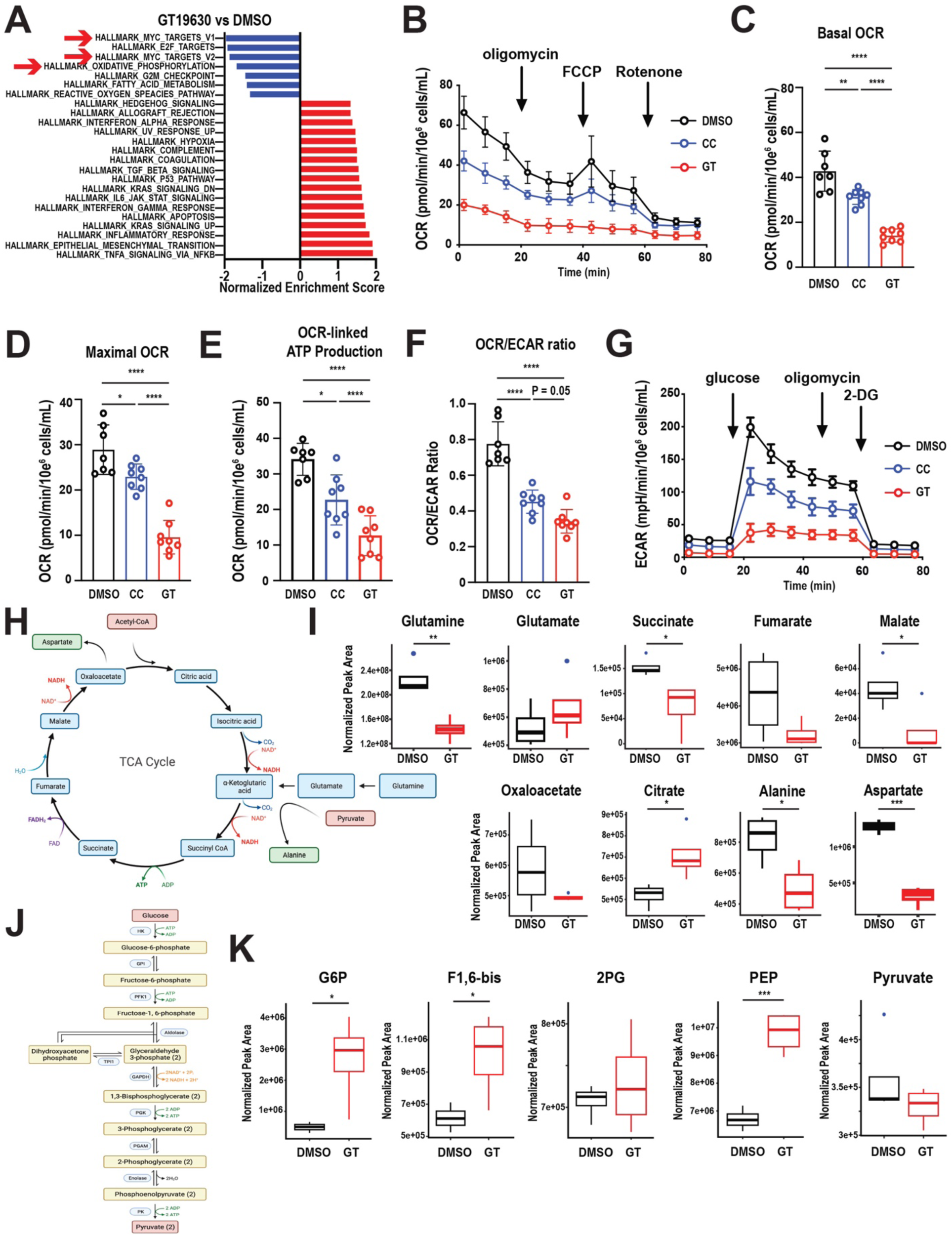
MYC/GSPT1 Degradation Induces Severe OXPHOS Inhibition. **A.** Normalized enrichment scores in significantly downregulated (blue) or upregulated (red) pathways from RNA-seq analyses of HL-60 GT19630-sensitive (GTS) cells treated with DMSO or 5 nM GT19630 for 12 hours. Downregulation of MYC targets and a pathway for oxidative phosphorylation are highlighted by red arrows. **B.** Oxygen consumption rate (OCR) measured by Seahorse analysis in HL-60 GTS cells treated with DMSO, 100 nM CC-90009 (CC) or 5 nM GT19630 (GT) for 12 hours. FCCP, carbonyl cyanide-p-trifluoromethoxyphenylhydrazone, **C-F.** Measures of basal OCR (**C**), maximal OCR (**D**), OCR-linked ATP production (**E**) and OCR/ECAR ratio (**F**) in HL-60 GTS cells treated with DMSO, 100 nM CC or 5 nM GT for 12 hours. **G.** Extracellular acidification rate (ECAR) determined by Seahorse analysis in HL-60 GTS cells treated with DMSO, 100 nM CC or 5 nM GT for 12 hours. 2-DG, 2-deoxy-D-glucose. **H.** Schema of the TCA cycle for metabolic flux analyses (MFAs). **I.** Levels of TCA cycle intermediates in HL-60 GTS cells treated with DMSO or 5 nM GT for 12 hours. **J.** Schema of glycolysis for MFA. **K.** Levels of glycolysis intermediates in HL-60 GTS cells treated with DMSO or 5 nM GT for 12 hours. Results are replicates of four independent measurements with mean +/- SD. Blue dots represent outliers for I and K. * *P* < 0.05, ** *P* < 0.01, *** *P* < 0.001, **** *P* < 0.0001.

### Dual MYC/GSPT1 degradation inhibits glutaminolysis and blocks the TCA cycle

To obtain further details of the metabolomic sequelae induced by MYC/GSPT1 degradation, we performed stable isotope-tracing metabolic flux analysis (MFA) using ^13^C_5_,^15^N_2_-labeled glutamine (**Fig. 3A**). First, we investigated the effects of GT19630 on glutamine utilization. GT19630 increased levels of non-labeled glutamate and reduced levels of newly incorporated ^13^C_5_^15^N_1_-labeled (m+5) glutamate. Subsequently, GT19630 reduced levels of glutamine, as well as levels of m+2 succinate, m+4 aspartate and m+2 citrate, while increasing the level of m+4 succinate, the direct intermediate produced after one passage through the TCA cycle (**Fig. 3B-E**). These findings, combined with the reduced glutamine levels (**Fig. 2I**), suggest that GT19630 inhibits the TCA cycle at the level of glutamate synthesis. ^13^C_4_-labeled (m+4) and m+2 citrate levels were increased and decreased, respectively, indicating reduced flux through the TCA cycle by GT19630 following citrate synthesis (**Fig. 3E**). To dissect the impact of GT19630 on glucose metabolism, we then focused on MFA using ^13^C_6_-labeled glucose (**Fig. 3F**).

**Fig. 3.**
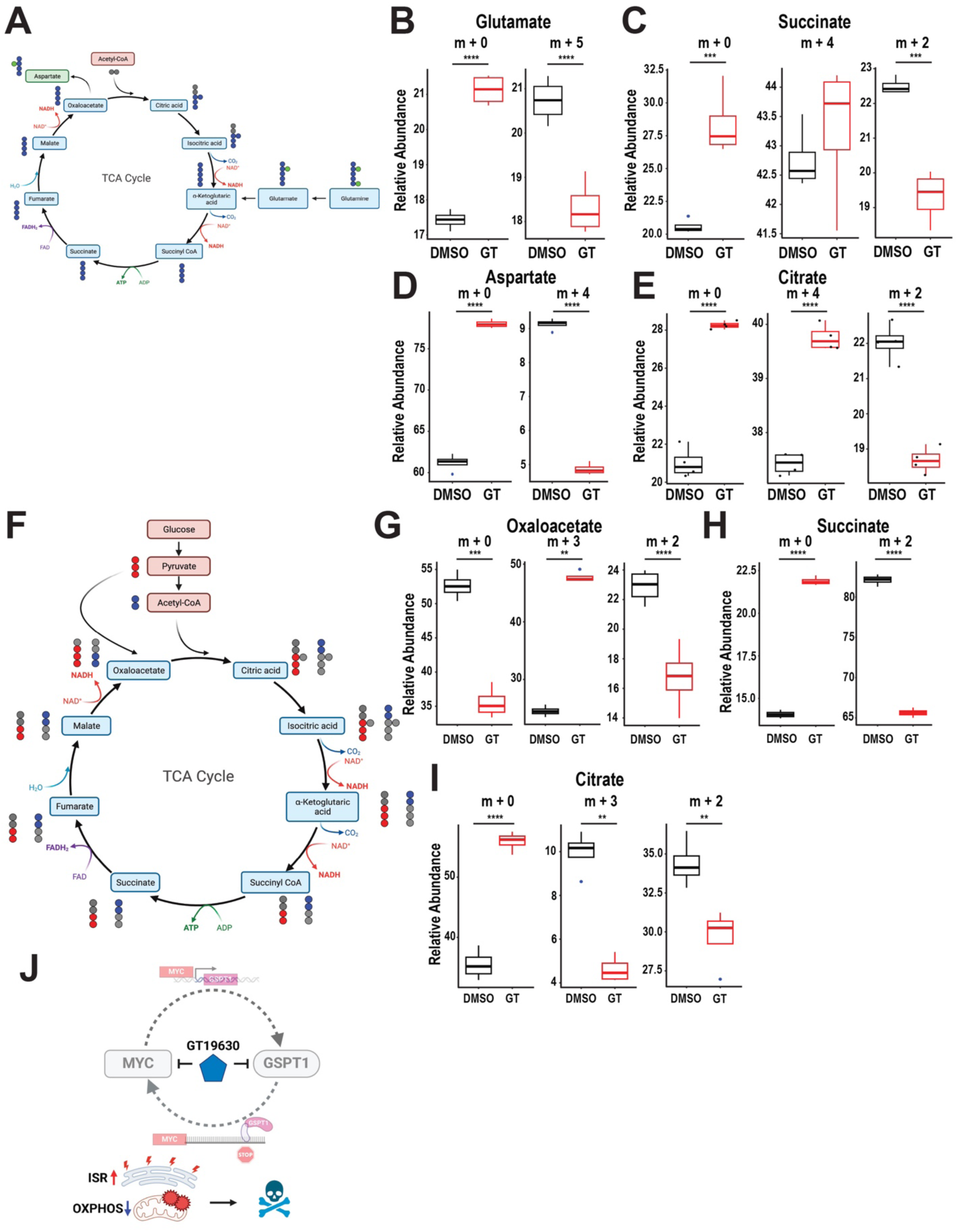
MYC/GSPT1 Degradation Inhibits TCA Cycle Flux. **A.** Map of TCA cycle for ^13^C_6_,^15^N_2_-glutamine–supplemented MFA. ^13^C_6_ carbon and ^15^N_2_ nitrogen are shown as blue and green dots, respectively. **B-E, G-I.** Levels of intermediates in HL-60 GTS cells treated with DMSO or 5 nM GT19630 (GT) for 12 hours. **B.** m+0 and m+5 glutamate levels. **C.** m+0, m+4 and m+2 succinate levels. **D.** m+0 and m+4 aspartate levels. **E.** m+0, m+4 and m+2 citrate levels. **F.** Map of TCA cycle for ^13^C_6_-glucose–supplemented MFA. Pyruvate- and acetyl CoA-derived ^13^C_6_ carbons are shown as red and blue circles, respectively. **G.** m+0, m+3 and m+2 oxaloacetate levels. **H.** m+0 and m+2 succinate levels. **I.** m+0, m+3 and m+2 citrate levels. **J.** Schematic diagram of the modes of action of GT19630. ISR, integrated stress response; OXPHOS, oxidative phosphorylation. Results are replicates of four independent measurements with mean +/- SD. Blue dots represent outliers for B-E and G-H. ** *P* < 0.01, *** *P* < 0.001, **** *P* < 0.0001.

Interestingly, the levels of ^13^C_3_-glucose-labeled (m+3) oxaloacetate, which is synthesized from m+3 pyruvate, increased while those of m+3 citrate and m+2 succinate decreased, suggesting that the TCA cycle decelerated at the point of citrate synthesis from oxaloacetate (**Fig. 3G-I**). We observed that the levels of ^13^C_6_-glucose-labeled (m+6) G6P and m+3 PEP increased, while those of m+3 pyruvate decreased, suggesting accumulation of glycolysis intermediates that were unable to proceed through the TCA cycle (**Supplementary Fig. S5A-D**). The overall levels of pyruvate did not change (**Fig. 2K**), and the levels of m+3 pyruvate decreased (**Supplementary Fig. S5D**). Interestingly, intracellular lactate levels increased while those of extracellular lactate decreased (**Supplementary Fig. S5E, F**), suggesting an underlying cellular effort upon GT19630-induced inhibition of the TCA cycle to re-incorporate lactate as another source of energy in response to the lack of NADH due to TCA cycle inhibition^28,29^. Taken together, these data show that dual MYC/GSPT1 degradation profoundly decelerates flux through the TCA cycle at the stages of glutamate and citrate synthesis, thereby abrogating mitochondrial respiration and inducing lethality in MYC-driven leukemias (**Fig. 3J**).

### Anti-tumor activity of GT19630 is agnostic of *TP53* status in a broad spectrum of blood cancers

We analyzed MYC expression levels of hematologic malignancies in the Munich Leukemia Laboratory (MLL) 5K dataset ^30^. As expected, B-cell non-Hodgkin and high-grade B-cell lymphomas expressed the highest MYC levels, followed by mantle cell lymphomas, marginal zone lymphomas and multiple myelomas (**Fig. 4A**). We tested GT19630 in a wide range of cancer cell lines, including blood cancers, small cell lung cancers (SCLCs), triple-negative breast cancers and gliomas. Notably, most blood cancer and SCLC cell lines responded to GT19630 with IC50s of below 100 nM, suggesting efficacy against those tumor types (**Fig. 4B**). GT19630 induced apoptosis as determined by an annexin V–DAPI assay, with IC50 values of ≤10 nM in most cell lines. Colony-forming unit assays of normal erythroid or myeloid progenitors revealed higher IC50 values (43 – 44 nM), suggesting a potential therapeutic window (**Fig. 4C, D, Supplementary Table S1**).

**Fig. 4.**
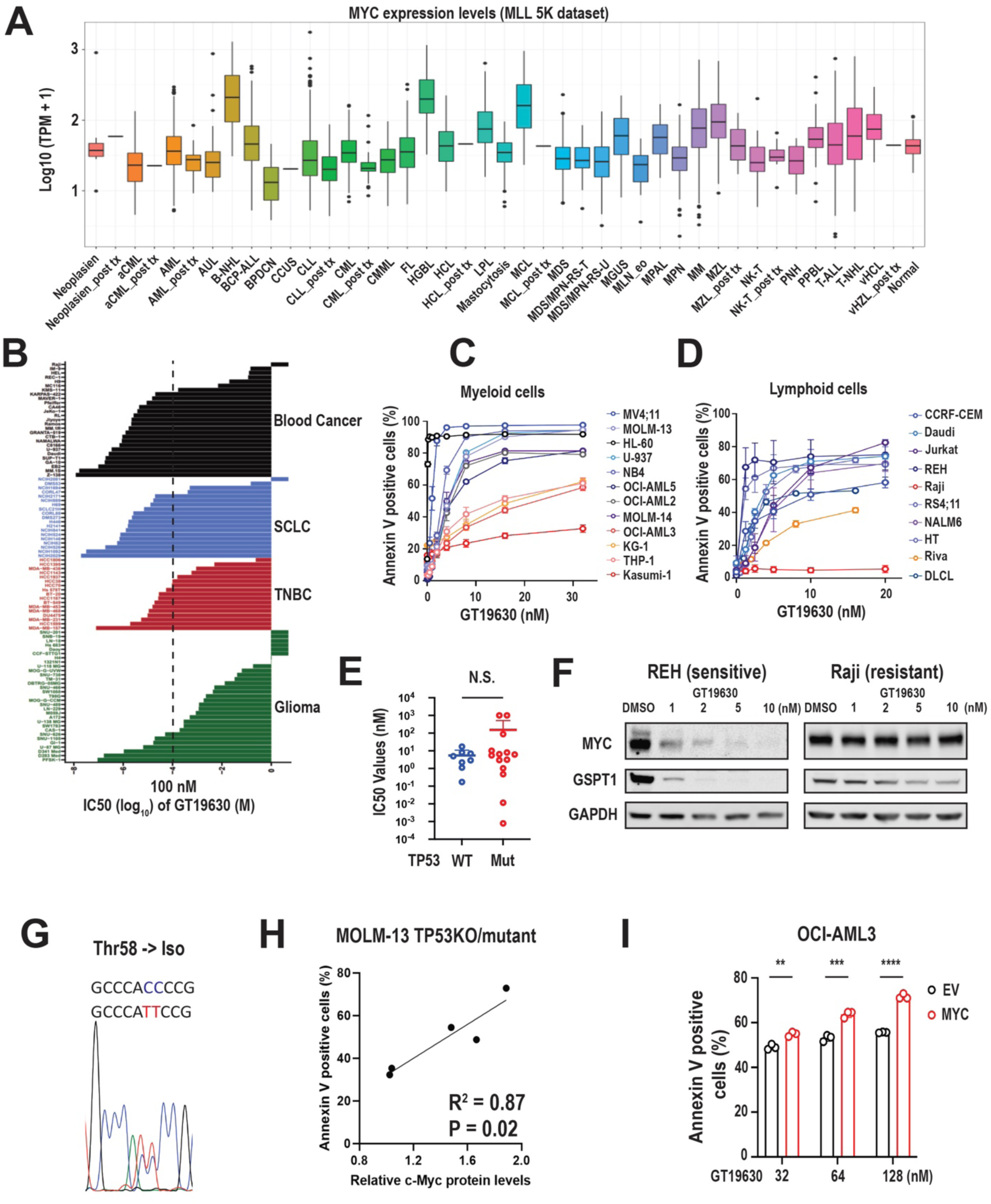
GT19630 induces TP53 independent cell death in hematologic malignancy cell lines with high expression of MYC. **A.** MYC expression levels among hematologic malignancies in the Munich Leukemia Laboratory 5K dataset. aCML, atypical chronic myeloid leukemia; AUL, acute undifferentiated leukemia; B-NHL, B-cell non-Hodgkin lymphoma; BCP-ALL, B-cell precursor acute lymphoblastic leukemia; BPDCN, blastic plasmacytoid dendritic cell neoplasm; CCUS, chronic cytopenia of unknown significance; CLL, chronic lymphocytic leukemia; CML, chronic myeloid leukemia; CMML, chronic myelomonocytic leukemia; FL, follicular lymphoma; HGBL, high-grade B-cell lymphoma; HCL, hairy cell leukemia; LPL, lymphoplasmacytic lymphoma; MCL, mantle cell lymphoma; MDS, myelodysplastic syndrome; MDS/MPN-RS-T, myelodysplastic syndrome/myeloproliferative neoplasm with ringed sideroblasts and thrombocytosis; MDS/MPN-RS-U, myelodysplastic syndrome/myeloproliferative neoplasm–unclassifiable; MGUS, monoclonal gammopathy of unknown significance; MLN-eo, myeloid or lymphoid neoplasm associated with eosinophilia; MPAL, mixed-phenotype acute leukemia; MPN, myeloproliferative neoplasm; MM, multiple myeloma; MZL, marginal-zone lymphoma; NK-T, natural killer/T-cell lymphoma; PNH, paroxysmal nocturnal hematuria; PPBL, persistent polyclonal B-cell lymphocytosis; T-ALL, T-cell acute lymphoblastic leukemia; T-NHL, T-cell non-Hodgkin lymphoma; vHCL, variant hairy cell leukemia. **B.** IC50 values after in vitro GT19630 treatment in panels of cancer cell lines that included blood cancers, small cell lung cancers (SCLCs), triple-negative breast cancer (TNBCs) and gliomas. The dashed line represents an IC50 of 100 nM. **C, D.** Percentages of annexin V– positive cells in myeloid leukemia (**C**) and lymphoid malignancy (**D**) cell lines treated with indicated concentrations of GT19630 for 48 hours. **E.** IC50 values of the cell lines from panels C and D classified by wild-type (WT) vs mutant (Mut) *TP53*. **F.** Western blot of MYC and GSPT1 in REH (GT19630-sensitive) and Raji (GT19630-resistant) cells. GAPDH served as loading control. **G.** Chromatogram of the Thr58 area of *MYC* in Raji cells. **H.** Correlation plot of baseline MYC protein levels and annexin V–positive cells in isogenic MOLM-13 cells with *TP53* knockout (KO) or mutations (R248Q, R175H, R273H and Y220C) treated with 40 nM GT19630 for 48 hours. **I.** Percentages of annexin V–positive cells in OCI-AML3 cells transfected with empty vector (EV) and MYC-overexpressing vector (MYC) and treated with indicated concentrations of GT19630 for 48 hours. N.S., not significant. ** *P* < 0.01, *** *P* < 0.001, **** *P* < 0.0001.

Importantly, the efficacy of GT19630 was *TP53* agnostic in both myeloid and lymphoid cell lines (**Fig. 4E**, **Supplementary Fig. S6A, B**). Surprisingly, while Daudi Burkitt’s Lymphoma cells were highly sensitive to GT19630 (as expected), Raji Burkitt’s Lymphoma cells were completely resistant. We observed GSPT1 but no MYC reduction in Raji cells compared to GT19630-sensitive REH cells (**Fig. 4F**). In Raji cells, we identified a *MYC* p.T58I mutation, which stabilizes MYC by preventing phosphorylation by GSK3b and subsequent polyubiquitination that leads to degradation ^31,32^ (**Fig. 4G**). Protein levels of MYC (**Fig. 4H**) but not of GSPT1 (**Supplementary Fig. S6C**) showed a significantly positive correlation with annexin V induction in MOLM-13 CRISPR-engineered cells carrying different *TP53* mutations ^33^. We treated OCI-AML3 cells overexpressing MYC with GT19630 (**Supplementary Fig. S6D-F**), and MYC overexpressing OCI-AML3 cells exhibited significantly greater apoptosis induction by GT19630 compared to control cells (**Fig. 4I**).

Taken together, our findings show that dual targeting of MYC/GSPT1 by protein degradation exerts high apoptogenic activity in blood cancer cells irrespective of *TP53* status, and that this effect is MYC dependent, and to a lesser degree, dependent on GSPT1.

### GT19630 exhibits anti-tumor activity in MYC-driven malignancies in vivo

Next, to investigate the anti-tumor efficacy of GT19630 in vivo, we first treated NOD/SCID mice injected with Daudi cells with vehicle or GT19630 (**Fig. 5A**). Compared to vehicle controls, GT19630 nearly completely eradicated circulating CD20-positive cells (**Fig. 5B**) and significantly prolonged mouse survival (**Fig. 5C**), with all mice treated with 3 mg/kg alive on day 42, compared with only 20% of vehicle-treated controls. To evaluate the efficacy and pharmacodynamic profiles of GT19630, BALB/c nude mice, subcutaneously inoculated with HL-60 cells, were treated with vehicle or GT19630 (**Fig. 5D**). Reduced tumor volumes (**Fig. 5E**) and reduced MYC protein levels (**Fig. 5F**) were observed in the GT19630-treated tumors compared to vehicle controls. The reduction of MYC was sustained for at least 24 and 48 hours after one injection of GT19630 (**Supplementary Fig. S7A**). Next, we injected mice subcutaneously with MM.1S cells and treated them with GT19630 or vehicle controls (**Supplementary Fig. S7B**). GT19630 significantly reduced MM.1.S tumor volumes at all dose levels compared to vehicle controls, even below baseline (**Supplementary Fig. S7C**).

**Fig. 5.**
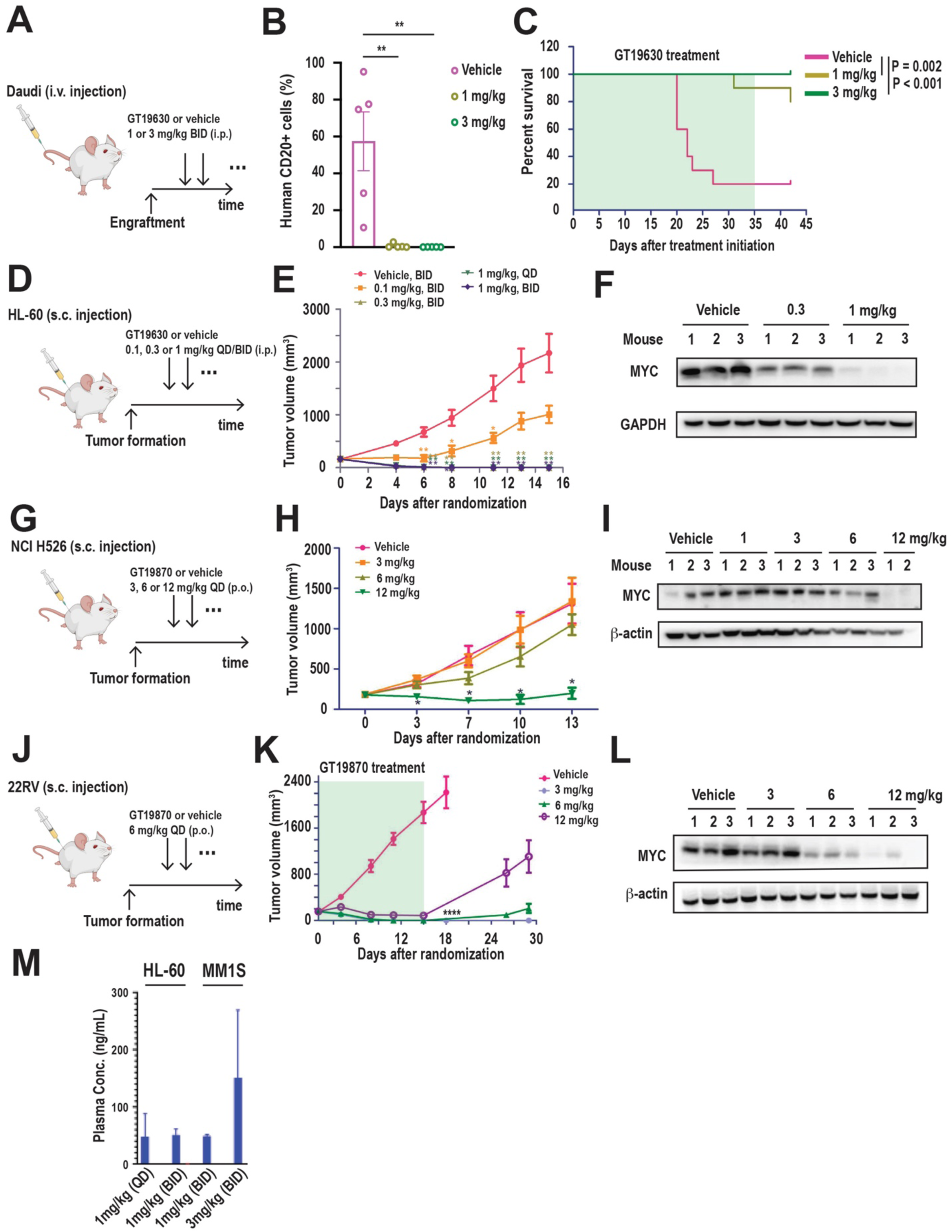
GT19630 exerts in vivo activity in MYC-driven hematologic tumor models with favorable in vivo pharmacokinetic and pharmacodynamic profiles. **A.** Schematic diagram of in vivo experiment. Mice were intravenously (i.v.) injected with Daudi cells into their tail veins and treated intraperitoneally (i.p.) with indicated doses of GT19630 or vehicle twice daily (BID). **B.** Percentages of human CD20-positive cells determined by flow cytometry in bone marrow samples from 5 different mice treated with vehicle or 1 mg/kg or 3 mg/kg of GT19630. **C.** Survival curves of mice treated with vehicle or 1 mg/kg or 3 mg/kg of GT19630. Treatment duration is shaded in green. **D.** Schematic diagram of in vivo experiment. Mice were subcutaneously (s.c.) injected with HL-60 cells and treated with indicated doses of GT19630 or vehicle once a day (QD) or BID. **E.** Size of HL-60 tumors in each treatment group at indicated days. **F.** Western blot of MYC in HL-60 tumors obtained from each treatment group (N = 3 each). GAPDH served as loading control. **G.** Schematic diagram of in vivo experiment. Mice were injected with NCI H526 cells and treated with GT19870 or vehicle with indicated doses every three days. **H.** Size of NCI H526 tumors in each treatment group at indicated days. **I.** Western blot of MYC in NCI H526 tumors obtained from each treatment group (N = 3 each, N = 2 for mice receiving 12 mg/kg). β-actin served as loading control. **J.** Schematic diagram of in vivo experiment. Mice were injected with 22RV1 cells and treated with GT19870 or vehicle with indicated doses every three days. **K.** Size of 22RV1 tumors in each treatment group at indicated days. **L.** Western blot of MYC in 22RV1 tumors obtained from each treatment group (N = 3 each). β-actin served as loading control. **M.** Plasma concentrations of GT19630 at 4 hours after the injections in the HL-60 and MM.1S tumor models. Mean values ± SEM were shown in all graphs in this figure.

To investigate the efficacy of MYC/GSPT1 protein degradation in MYC-driven small solid tumors in vivo, we developed GT19870, an oral formulation of GT19630 and tested it in small cell lung cancer (SCLC) and castration-resistant prostate cancer (CRPC) models. GT19870 significantly reduced volume of NCI H526 tumors (**Fig. 5G, H**), and reduced MYC protein levels (**Fig. 5I**). Similarly, GT19870 reduced the volume of 22RV tumors (**Fig. 5J, K**), and MYC protein levels (**Fig. 5L**). Lastly, to assess pharmacokinetic profiles of GT19630, we determined plasma concentrations of GT19630 in the HL-60 and MM.1.S models. One or two injections of 1 mg/kg or 3 mg/kg GT19630 yielded plasma concentrations of 50–150 ng/mL, corresponding to 77–231 nM GT19630 with a half-life of 4 – 8 hours (data not shown), which is substantially higher than the IC50 values of those cells (**Fig. 5M**). Taking these data together, MYC/GSPT1 protein degradation exerted promising anti-tumor efficacy in liquid and solid tumor models in vivo with favorable pharmacokinetic and pharmacodynamic profiles.

### *MYC* is overexpressed in immature CD34+ AML blasts, and dual MYC/GSPT1 degradation targets immature CD34+ AML cells

*MYC* overexpression is known to have leukemia-initiating potential in AML and is associated with worse survival ^34–36^; however, detailed profiling of MYC expression levels in different AML subtypes has not been performed. We analyzed The Cancer Genome Atlas dataset and found a relatively infrequent incidence of *MYC* amplifications or mutations in AML compared to other cancer types (**Fig. 6A**). However, MYC expression levels in AML were the third highest among different cancer types (**Fig. 6B**). By multivariate analyses of the MLL AML dataset we determined that AML with trisomy of chromosome 8, where *MYC* is located, exhibited a higher proportion of samples with high MYC expression (MYC^high^), as expected, followed by AMLs with *TP53* mutations and t(15;17) translocations, while those with inv.3, *FLT3* and *NPM1* mutations exhibited lower numbers of MYC^high^ and higher numbers of MYC^low^ samples (**Fig. 6C**). These results are consistent with the notion that MYC is regulated by *TP53* or *PML-RARa* ^37–39^.

**Fig. 6.**
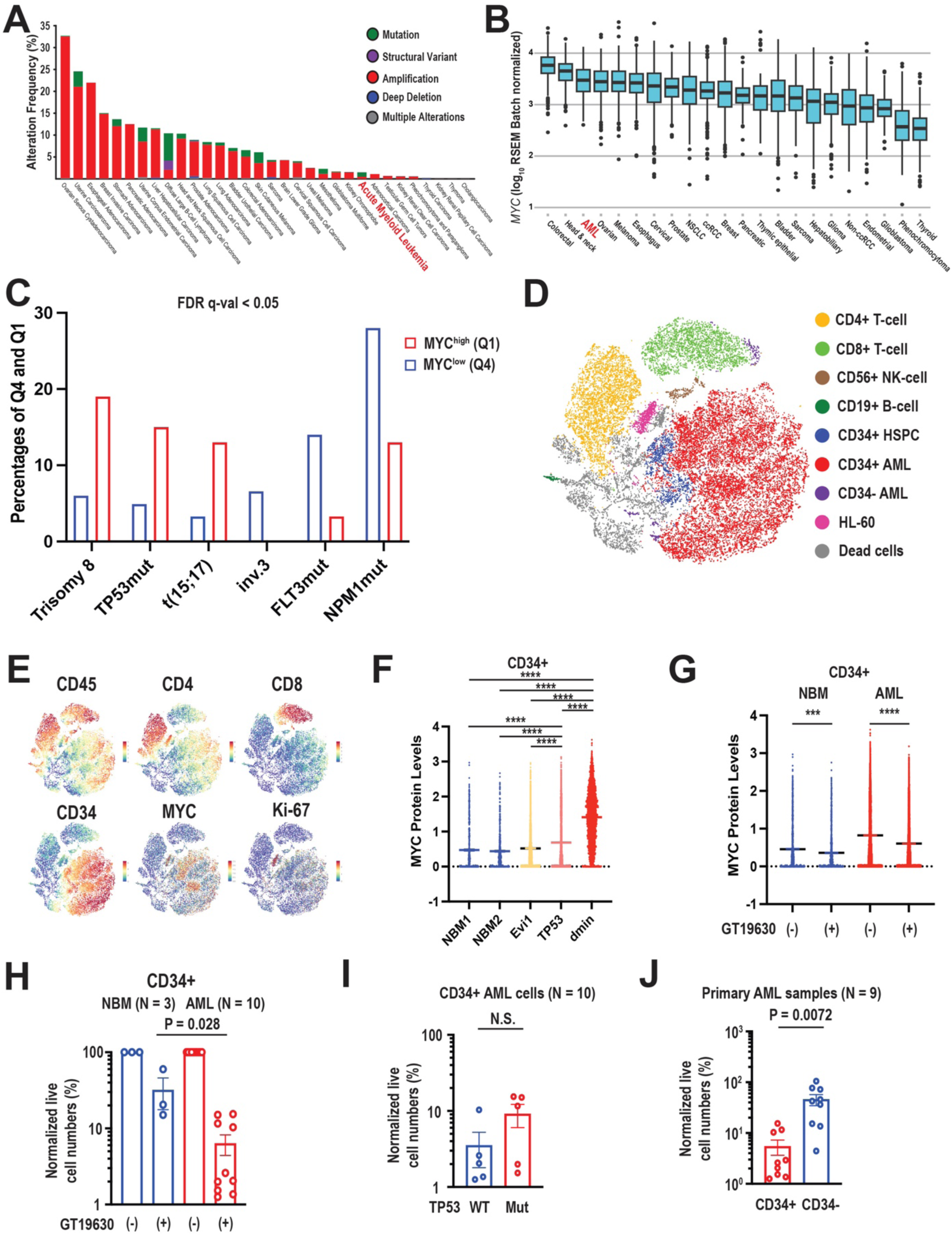
MYC/GSPT1 degradation targets immature CD34+ AML cells while sparing normal CD34+ hematopoietic stem/progenitor cells. **A.** Frequencies of *MYC* amplifications among 32 cancer types, including acute myeloid leukemia (AML) (highlighted in red), in the TCGA dataset. **B.** MYC expression levels of 22 cancer types, including AML (highlighted in red), in the TCGA dataset. **C.** Percentages of MYC^high^ (quartile 1) and MYC^low^ (quartile 4) expression among different subtypes of AML in the Munich Leukemia Laboratory dataset. Only subtypes that had statistical significance (false discovery rate [FDR] q-val < 0.05) are shown. **D.** Unsupervised clustering for indicated cell annotations of two normal bone marrow (NBM) and three AML samples with double-minute (dmin) chromosomes, *TP53* mutation (TP53) and 3q26 abnormality (Evi-1) subjected to single-cell mass cytometry (CyTOF). **E.** Protein levels of CD45, CD4, CD8, CD34, MYC and Ki-67 determined by CyTOF and unsupervised clustering as shown in panel D. **F.** MYC protein levels in the NBM and AML samples described in panel D. **G.** MYC protein levels in the NBM and AML samples before (-) and after (+) GT19630 treatment (64 nM) for 24 hours. **H.** Normalized live cell numbers of NBM and AML samples before (-) and after (+)GT19630 treatment (64 nM) for 72 hours. **I.** Normalized live cell numbers of TP53 wild-type (WT) and mutant (Mut) AML samples treated with 64 nM GT19630 for 72 hours. **J.** Normalized live cell numbers of CD34-positive and -negative fractions in AML samples treated with 64 nM GT19630 for 72 hours. Paired *t* test was used for statistical comparison. N.S., not significant. *** *P* < 0.001, **** *P* < 0.0001.

To evaluate protein levels of MYC in normal hematopoietic stem/progenitor cells (HSPCs) and AML leukemia stem/progenitor cells (LSPCs), we performed single-cell mass cytometry (CyTOF) of normal bone marrows (NBMs) and AML samples with HL-60 cells as positive control. CyTOF analyses identified 25 distinct clusters (**Supplementary Fig. S8A**), and we annotated those based on surface marker expression patterns (**Fig. 6D**). HL-60 cells, serving as positive control for high MYC levels, were characterized by high MYC, CLL-1 and Ki-67 protein levels (**Fig. 6D, E, Supplementary Fig. S8B**) ^40^. NBM HPSCs and AML LSPCs are shown in blue and red clusters, respectively, based on CD34 expression levels. Compared with NBM HSPCs, AML LSPCs from patients carrying the double-minute chromosome abnormality or *TP53* mutations exhibited significantly higher MYC protein levels (**Fig. 6F**). GT19630 treatment reduced MYC protein levels in both NBM HSPCs and AML LSPCs, with greater reduction in AML LSPCs (**Fig. 6G**).

We next treated ten primary AML samples (characteristics shown in **Supplementary Table S2**) with GT19630 and observed dose-dependent cell death (**Supplementary Fig. S8C, D**). Notably, AML LSPCs were significantly more sensitive to GT19630 than NBM HSPCs, with approximately one log_10_ difference in cell death induction (**Fig. 6H**), suggesting a therapeutic window between normal HSPCs and AML LSPCs. *TP53* mutant AML CD34+ cells were as sensitive to GT19630 as *TP53* wild-type AML CD34+ cells (**Fig. 6I**). AML LSPCs were more sensitive to GT19630 than CD34-negative AML cells (**Fig. 6J**). In summary, our findings show that *MYC* and MYC protein levels are higher in AML LSPCs than in normal HSPCs, resulting in a therapeutic window.

### Dual MYC/GSPT1 degradation results in profound anti-leukemia activity in AML with tolerable toxicity profiles in vivo

Given the promising efficacy of GT19630 in the HL-60 AML model in vivo, we next evaluated the efficacy of GT19630 in an AML patient-derived xenograft (PDX) model (**Fig. 7A**). We treated NOD/SCID mice injected with PDX AML cells carrying *FLT3-ITD*, *IDH2* p.R140Q and *CEBPA* mutations established from a patient whose AML had relapsed after induction chemotherapy. GT19630 eradicated AML cells in circulation, bone marrows and spleens, resulting in significantly reduced spleen weights compared to vehicle treatment (**Fig. 7B, C, Supplementary Fig. S9A, B**). Previously, we reported that AML cells refractory to venetoclax overexpress MYC^10^. We also observed increased MYC and GSPT1 protein levels in MV4;11 venetoclax-resistant (Ven-R) compared to venetoclax-sensitive (Ven-S) cells (**Fig. 7D**). GT19630 reduced levels of MYC, GSPT1 and intriguingly, MCL-1, a major resistance factor to Ven^41^, in MV4;11 Ven-R cells (**Fig. 7E**). To further validate the efficacy of GT19630 in Ven-R AML in vivo, we injected GFP/luciferase-transduced MV4;11 Ven-S and Ven-R cells into NSG mice (**Fig. 7F**). GT19630 significantly reduced leukemia burden (**Fig. 7G, H**) and profoundly reduced both Ven-S and Ven-R cells in bone marrows and spleens (**Supplementary Fig. S9C-F**). GT19630 greatly improved the survival of both MV4;11 Ven-S and Ven-R groups, resulting in >300% prolongation of the Ven-R group (**Fig. 7I**). We determined tumor burden by BLI in the remaining mice on day 175 and found comparable levels in the surviving mice treated with GT19630 and NSG mice not injected with AML cells (**Supplementary Fig. S9G**). We then discontinued GT19630 and monitored the remaining mice. There were no leukemia-related deaths from day 175 through termination of the experiment at day 294, suggesting that GT19630 potentially eradicated MV4;11 Ven-R and Ven-S cells. These data suggest that GT19630 has promising anti-leukemia efficacy in both, Ven-S and Ven-R AML in vivo.

**Fig. 7.**
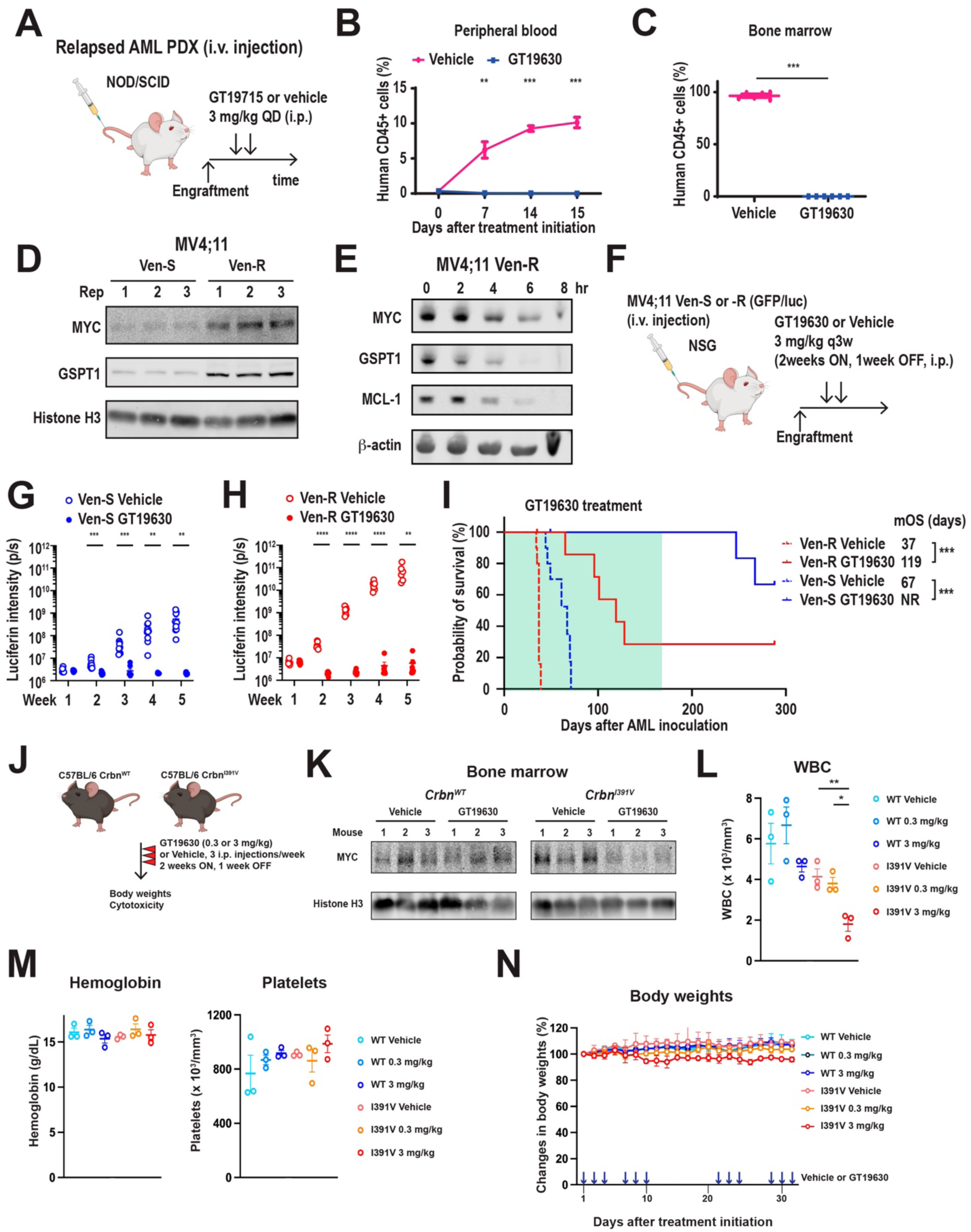
MYC/GSPT1 degradation exhibits activities in AML in vivo. **A.** Schematic diagram of in vivo experiment using NOD/SCID mice injected with AML patient-derived xenograft (PDX) cells and treated with vehicle or GT19630. **B.** Percentages of circulating human CD45-positive cells in mice treated with vehicle or GT19630. **C.** Percentages of human CD45-positive cells in bone marrow samples in mice treated with vehicle or GT19630. **D.** Protein levels of MYC and GSPT1 in venetoclax-sensitive (Ven-S) and -resistant (Ven-R) MV4;11 cells. Histone H3 served as loading control. **E.** Schematic diagram of in vivo experiment using NSG mice injected with green fluorescent protein [GFP]/luciferin–labeled MV4;11 Ven-S and Ven-R cells and then treated as indicated with vehicle or GT19630. Every-3-week (q3w) cycles of 2 weeks on and 1 week off were used. **F, G.** Luciferin intensities of mice with MV4;11 Ven-S (**F**) or Ven-R (**G**) cells treated with vehicle or GT19630. **H.** Survival curves of NSG mice injected with Ven-S or -R cells treated with vehicle or GT19630. The GT19630 treatment period is shaded in green. mOS, median overall survival.

**Fig. 8.**
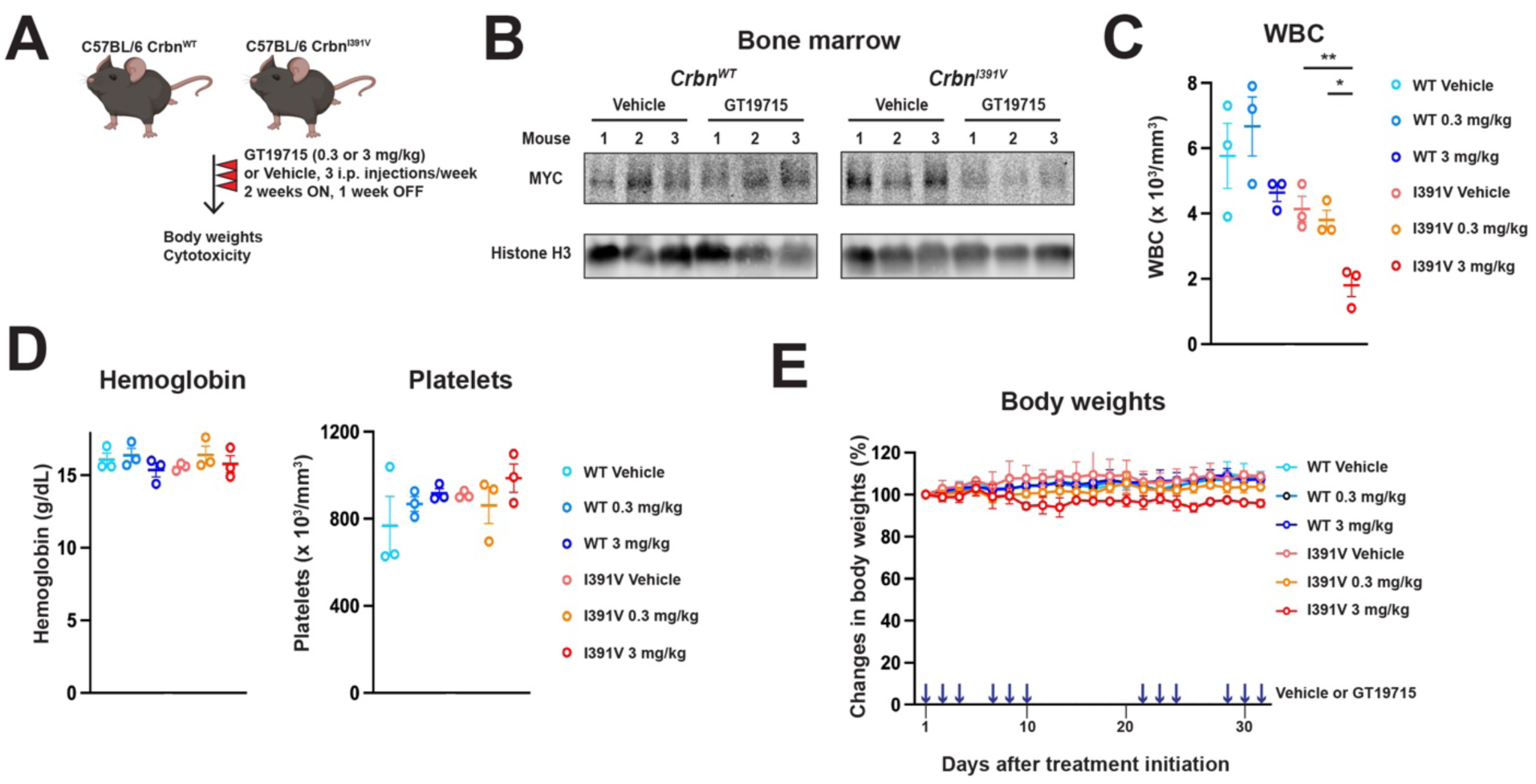
GT19175 is tolerated in humanized *Crbn^I391V^* mice. **A.** Schematic diagram of in vivo experiment using C57BL/6 mice with wild-type and I391V mutant *Crbn* (*Crbn^WT^*, *Crbn^I391V^*) treated as indicated with vehicle or GT19630. **B.** MYC protein levels in bone marrow samples from *Crbn^WT^* and *Crbn^I391V^* mice treated with vehicle or GT19630 (3 mg/kg). Histone H3 served as loading control. **C.** White blood cell (WBC) counts in peripheral blood samples (N = 3 in each group) in *Crbn^WT^* and *Crbn^I391V^* mice with indicated treatments. **D.** Hemoglobin and platelet levels in peripheral blood samples (N = 3 in each group) in *Crbn^WT^*and *Crbn^I391V^* mice with indicated treatments. **E.** Changes in body weights of *Crbn^WT^* and *Crbn^I391V^* mice during the indicated treatments. Vehicle or GT19630 treatments are indicated by blue arrows.

Murine Crbn is insensitive to degrader utilizing human CRBN^42^. We therefore treated C57BL/6 mice carrying *Crbn^WT^*and the *Crbn^I391V^* mutation, which renders *Crbn^I391V^*mice sensitive to Crbn-mediated protein degradation^42^, with the same GT19630 dosing schedule used in the leukemia models (**Fig. 7J**). Indeed, MYC protein levels decreased in bone marrow cells of *Crbn^I391V^*but not *Crbn^WT^* mice (**Fig. 7K**). Consistently, MYC protein levels decreased in granulocytes (Supplementary Fig. 9A) and GSPT1 protein were decreased in lymphocytes (Supplementary Fig. 9B) in *Crbn^I391V^* but not in *Crbn^WT^* mice. We observed decreased numbers of white blood cells (WBCs) in *Crbn^I391V^* mice treated with 3 mg/kg GT19630 compared to vehicle or 0.3 mg/kg GT19630 (**Fig. 7L**). GT19630 did not induce reductions in hemoglobin and platelet levels (**Fig. 7M**). Murine bone marrow CD45-positive cells increased significantly at the end of GT19630 treatment in *Crbn^I391V^*mice, potentially suggesting recovery from myelosuppression (Supplementary Fig. S9C). GT19630 only moderately decreased the body weights of *Crbn^I391V^*mice by around 5% at most during treatment (**Fig. 7N**). No mice died and no obvious adverse events were observed. Taken together, these data show that dual MYC/GSPT1 protein degradation induces minimal, reversible myelosuppression in mice carrying humanized *Crbn*.

## Discussion

MYC, since its discovery in Burkitt’s Lymphomas carrying the *IgH/MYC* translocation, has been a challenging therapeutic target. MYC family proteins N-MYC and MYCL share structural similarities in MYC boxes and the bHLH-LZ domain and are highly expressed in hematologic and solid tumors, including SCLCs and high-grade gliomas^3^. Numerous attempts have been made to target MYC:MAX interaction or to indirectly target MYC by bromodomain inhibition^43^. Here we report, for the first time, the direct degradation of MYC.

The translational control by MYC has been investigated in depth, with focus on ribosome biogenesis, translation initiation and elongation^44^. eIF4E, the rate-limiting factor of translation initiation, is one of MYC’s targets, and overexpression of eIF4F complex facilitates cancer initiation^45^. Indeed, we previously reported targeting eIF4A, the enzymatic core of the complex, with rohinitib as an effective therapeutic approach for AML^46^. However, the role of MYC in translation termination is not fully established. Here, we report that MYC binds to the TSS of *GSPT1* and transcriptionally regulates the gene, a decoder for stop codons, through GTP hydrolysis in cooperation with eRF1, facilitating translation termination^14^. Importantly, our SCR reporter system for MYC revealed that GSPT1 promotes translation termination of *MYC* mRNA and that degradation of GSPT1 induces SCR of MYC, uncovering a novel, co-regulatory positive feedback loop between MYC and GSPT1 that was discovered to be “druggable” with GT19630.

We observed promising anti-tumor activities by targeting MYC and GSPT1 proteins with GT19630 in different models of MYC-driven hematologic malignancies, including Burkitt’s Lymphoma, multiple myeloma, AML, SCLC and CRPC. GT19630 binds to both MYC and GSPT1 and induces much enhanced apoptosis compared to selective GSPT1 degradation. Although disease outcomes of Burkitt’s Lymphoma, the quintessential MYC-driven hematologic malignancy, have improved since the introduction of rituximab^47^, relapsed cases are highly drug resistant, and newly diagnosed patients with elevated lactate dehydrogenase or involvement of the central nervous system have aggressive disease ^48^. Patients with multiple myelomas carrying *MYC* rearrangements also have unfavorable survival ^49^. In diffuse large B-cell lymphomas, high expression of MYC target genes was identified as a high-risk signature, and MYC was ranked as the top essential gene by genome-wide CRISPR screening^50^. Double- and triple-hit diffuse large B-cell lymphomas are particularly challenging MYC-driven hematologic malignancies despite recent advances in CAR-T cell therapies^51,52^. We recently reported that AML cells from patients with acquired resistance to venetoclax-based therapies, who have a median survival of only 2.4 months^53^, expressed higher MYC protein levels compared to cells from patients with AML sensitive to venetoclax, and that knockdown of *MYC* restores venetoclax sensitivity^10^. We and others also found that *TP53*-mutant AMLs, the type of AML with the shortest survival, express elevated *MYC* levels, based on the counter-regulatory roles of *TP53* and *MYC*^54,55^. Thus, targeting MYC is of particular relevance to improve outcomes in patients with MYC-driven hematologic malignancies, including TP53 mutant malignancies for which still no effective therapeutic options exist. In the current study, GT19630 exerted profound reductions of disease burden and prolongation of survival in numerous types of hematologic malignancies in vitro and in Burkitt’s Lymphoma, multiple myeloma, Ven-R AML and chemotherapy-resistant AML PDX models in vivo. Further, oral GT19870 treatment significantly reduced tumor burden in SCLC and CRPC. Data suggest MYC/GSPT1 protein degradation as a promising novel treatment modality. We have demonstrated high activities against MYC expressing tumors while a mutation at T58 in MYC in Raji cells abrogated its activity.

Targeting of MYC may raise concerns regarding its toxicity, considering that MYC controls homeostasis in normal cells and tissues. A recent phase 1 clinical trial that investigated the safety of OMO-103, a MYC:MAX interaction–inhibiting miniprotein, in patients with advanced solid tumors demonstrated tolerability of OMO-103 with favorable toxicity profiles^56^. We determined a preclinical therapeutic differential between AML LSPCs and normal HSPCs of approximately 1 log_10_ difference in sensitivity. GT19630 reduced MYC and GSPT1 protein levels in hematopoietic cells but induced only moderate leukopenia in humanized *Crbn^I391V^* mice. Post-natal Myc loss is tolerated in mice^57^, supporting our findings. However, these need to be extended by additional toxicity studies in other species. GSPT1 degraders have been investigated in preclinical and clinical studies, with lack of MYC reduction in vivo^17,58,59^.

MYC is involved in a plethora of metabolic steps in non-malignant and cancer cells^19^. To meet the increased biosynthetic demand of nutrients and metabolites for rapid proliferation, cancer and leukemia cells rely on *MYC* through aerobic glycolysis (the Warburg effect) and glutaminolysis^19^. Our initial hypothesis assumed that MYC/GSPT1 degradation would induce profound inhibition in glycolysis, based on the notion that MYC controls virtually all genes involved in glycolysis^20^. However, RNA-seq and MFA revealed that dual MYC/GSPT1 degradation by GT19630 primarily induced profound inhibition of OXPHOS and disruption of the TCA cycle at the glutamate and citrate synthesis levels, while glycolysis intermediates accumulated as a consequence of the TCA cycle perturbation in MYC-driven HL-60 cells. This finding is of particular relevance in AML as leukemia stem cells in AML rely on OXPHOS while normal HSPCs utilize glycolysis^60,61^, which provides a reason of the therapeutic window observed between CD34+ AML LSPCs and NBM HSPCs, in addition to the observed difference in MYC protein levels. While proliferating leukemia stem/progenitor cells (LSPCs) depend on MYC, quiescent LSPCs are not but are highly sensitive to BCL-2 inhibition ^62^, resulting in profound reduction of LSPCs^63^. Another metabolic aspect of GT19630’s action is its downregulation of fatty acid metabolism. Levels of total and m+4 citrate were increased after GT19630, while levels of m+2 citrate, which is produced after one passage of the TCA cycle, were decreased. These findings potentially suggest that citrate consumption for fatty acid biosynthesis is also abrogated by GT19630, providing intriguing topics for further investigation.

The impact of dual MYC/GSPT1 degradation on immune/tumor interaction and the immune microenvironment also merits future investigation. MYC has been recognized as a global regulator of the innate and adaptive immune response^64,65^. Our RNA-seq data showed upregulation of a multitude of immune reactive pathways in HL-60 GTS cells after GT19630, requiring further exploration. Furthermore, mechanisms of resistance to MYC/GSPT1 degradation are yet not understood. We observed reduced expression levels of CRBN in HL-60 GTR cells (data not shown), partially explaining resistance to GT19630 as observed in other studies in myeloma models and patients resistant to immunomodulatory agents or CELMoDs^66,67^. As noted, we observed only minimal toxicity in *Crbn^I391V^* mice treated with GT19630, but further toxicology studies in larger animals and feasibility studies in humans are needed. These aspects are limitations of the current study.

In conclusion, the novel co-regulatory feedback loop of MYC and GSPT1 has been established as an actionable target of protein degradation, with promising anti-cancer efficacy in multiple models of MYC-driven malignancies. Further investigations to translate these findings into the clinic are warranted.

## Acknowledgments

We appreciate Research Medical Library, MD Anderson Cancer Center for providing editorial assistance. We used Biorender to create schematic figures.

## Author contributions

Conceptualization: Y.N., L.M. and M.A. Methodology: Y.N., E.S., V.I. and S.T. Investigation: Y.N., V.I., N.B., E.A., D.S., H.M., L.B.O, S.K., S.S., A.B., Y.T., Q.Z., Z.Y., H.Y. and D.C. Visualization: E.A., E.S. and T.T. Funding acquisition: Y.N., N.B., S.T. and M.A. Project administration: P.Y.M., B.Z.C., A.M., K.S., S.B. and T.H. Supervision: S.T. and M.A. Writing (original draft): Y.N. Writing (reviews and edits): M.A.

## Data Access Statement

All of the materials used in the study, except reagents obtained under the material transfer agreement between the University of Texas MD Anderson Cancer Center and pharmaceutical companies, will be provided upon appropriate requests (contact). RNA-seq datasets will be available on a publicly available repository upon acceptance. All data needed to evaluate the conclusions in the paper are present in the paper and/or the Supplementary Materials.

## Ethics Statement

All patient samples were obtained and used under the Institutional Review Board Protocol LAB02-395 approved by the University of Texas MD Anderson Cancer Center Institutional Review Board. Animal experiments were performed under the IACUC protocol 00001774-RN02. Detailed information is described in the Supplementary Materials and Methods.

## Materials and methods

### Primary AML samples

All primary AML samples from patients and normal bone marrow mononuclear cells from healthy donors were processed as described previously^1^. All samples were collected with consent by patients under the protocols LAB01-473 and LAB02-395 approved by the institutional review board of The University of Texas MD Anderson Cancer Center. Briefly, freshly obtained samples were isolated by density-gradient centrifugation using Lymphocyte Separation Medium (Corning) and cryopreserved. Isolated cells were subjected to red blood cell lysis using BD Pharm Lyse lysing solution (BD Biosciences) and maintained in RPMI 1640 medium supplemented with 20% fetal bovine serum (FBS) with co-culturing with human mesenchymal stromal cells (cell density of 6,000/cm^2^) for indicated treatment durations.

### Animal experiments

#### Daudi systemic lymphoma model

Daudi cells were cultured in RPMI 1640 medium containing 20% FBS for 3-4 days. The cell density was adjusted to 5 x 10^7^ cells /mL with pre-cooled phosphate-buffered saline (PBS). NOD/SCID mice (n = 20; Beijing Vital River Laboratory Animal Technology) received tail vein injection of 5 x 10^6^ cells suspended in 100 µL PBS/mouse. Three days after inoculation, mice were randomized according to body weight; 15 mice were selected and grouped into three groups with 5 mice in each group. GT19630 was dissolved into 5% ethanol, 5% solutol HS15 in PBS (vehicle) and was administered by intraperitoneal (i.p.) injections with doses of 0.1 mg/kg and 0.3 mg/kg every 3 days. At the end of the experiment, mice were euthanized, bone marrow cells were harvested and 1 x 10^6^ cells were subjected to CD20 staining for 30 min. Stained samples were subjected to flow cytometry analyses.

#### HL-60 tumor xenograft model

HL-60 cells were cultured in RPMI 1640 medium containing 20% FBS for 3–4 days. The density of HL-60 cells was adjusted to 5 x 10^7^ cells/mL with a pre-cooled PBS and Matrigel (1:1) mixture. BALB/c nude mice (n = 40; Beijing Vital River Laboratory Animal Technology) were inoculated with 1 x 10^7^ cells/200 µL subcutaneously in the back fat pad on the right side. Tumor growth was observed daily after inoculating HL-60 cells. When the mean volume of tumor reached the range of 100–200 mm^3^, mice were randomized according to tumor volume into five groups with 8 mice/group. Vehicle and GT19630 were formulated as described above and GT19630 was administered with dosing of 0.1 mg/kg (twice a day), 0.3 mg/kg (twice a day), 1 mg/kg (once a day) or 1 mg/kg (twice a day) by i.p. injection every 3 days. Tumor sizes were determined every 2–3 days starting on day 4 after treatment initiation.

#### MM.1S tumor xenograft model

MM.1S cells were cultured in RPMI 1640 medium containing 20% FBS for 3–4 days. The density of cells was adjusted to 2.5 x 10^7^ cells/mL with pre-cooled PBS and Matrigel (1:1) mixture. NOD/SCID mice (n = 24; Beijing Vital River Laboratory Animal Technology) were inoculated with 5 x 10^6^ cells/mouse in 200 µL PBS subcutaneously on the right side back of fat pad. Tumor growth was observed daily after inoculating MM.1S cells. When the mean volume of tumor reached 150–200 mm^3^, mice were randomized according to tumor volume into four groups with 6 mice/group. Vehicle and GT19630 were formulated as described above and GT19630 was administered with dosing of 0.3 mg/kg (twice a day), 1 mg/kg (twice a day) or 3 mg/kg (twice a day) by i.p. injection every 3 days. Tumor sizes were determined every 3–4 days starting on day 3 after treatment initiation.

#### NCI H526 model

NCI H526 small cell lung cancer cells were cultured in RPMI 1640 medium containing 20% FBS and 1% penicillin/streptomycin. On the day of injection, cells were digested by trypsin and injected into female BALB/C nude mice (4-6 week old, n = 24; Beijing Vital River Laboratory Animal Technology) with 5 x 10^6^ cells/mouse in 200 µL PBS subcutaneously on the right side back of fat pad. When the mean volume of tumor reached 150–250 mm^3^, mice were randomized according to tumor volume into four groups with 6 mice/group. Vehicle (5% DMSO, 5% solutol HS15 in PBS) or GT19870 dissolved in vehicle solution with doses of 3, 6 or 12 mg/kg were administered by oral gavage. Tumor sizes were measured twice a week.

#### 22RV1 model

22RV1 prostate cancer cells were cultured in RPMI 1640 medium containing 20% FBS and 1% penicillin/streptomycin. On the day of injection, cells were digested using trypsin, and BALB/C nude mice (4-6 week-old, were inoculated with 5 x 106 cells/mouse in 200 µL PBS subcutaneously on the right side back of fat pad. When the mean volume of tumor reached 150–250 mm^3^, mice were randomized according to tumor volume into four groups with 6 mice/group. Vehicle or GT19870 were administered and mice were monitored in the same way as NCI H526 model.

#### Patient-derived xenograft (PDX) AML model

PDX AML AM7577 cells were injected with a density of 2 x 10^6^ cells/100 µL into NOD/SCID mice (n = 12; Shanghai Lingchang Biotechnology). Engraftment of AML cells was confirmed by the presence of 0.5–2% of human CD45+ (hCD45+) cells in peripheral blood specimens. Mice were randomized according to percentages of hCD45+ cells into two groups. Vehicle was formulated with 5% DMSO, 5% solutol HS15 in PBS, and GT19630 was dissolved in the vehicle. GT19630 was administered at a dose of 3 mg/kg (once a day). After randomization, percentages of hCD45+ cells were determined in peripheral blood samples every week. Since all mice receiving vehicle became moribund on day 15, all mice were euthanized and bone marrow cells were harvested on that day to compare percentages of hCD45+ cells between the vehicle and GT19630 groups.

#### MV4;11 venetoclax-sensitive (Ven-S) and -resistant (Ven-R) AML models

MV4;11 Ven-S and Ven-R cells labeled with green fluorescent protein (GFP)/luciferase were developed through chronic exposure to <u>venetoclax</u> as described previously^2^. MV4;11 Ven-S and Ven-R cells were cultured in RPMI 1640 media supplemented with 10% FBS. NSG mice (n = 40; The Jackson Laboratory) were injected with 0.5 x 10^6^ cells in 100 µL PBS/mouse. Engraftment was confirmed by positive signals of luciferin determined by bioluminescence imaging using an IVIS-200 imager. On day 6 after inoculation with MV4;11 Ven-S and Ven-R cells, mice were randomized according to luciferin intensities into two groups. Vehicle and GT19630 were prepared as described above and administered with dosing of 3 mg/kg (once a day) by i.p. injection three times a week; one cycle of treatment consisted of injections in the first and second week and no injection in the third week. Treatment was discontinued at day 175.

#### Humanized *Crbn^I391V^*mice

Wild-type C57BL/6 mice (n = 9) and *Crbn^I391V^* (C57BL/6-*Crbn^tm1.1Ble^*/J) mice (n = 9) were purchased from The Jackson Laboratory and randomized in three groups based on body weights. Vehicle and GT19630 were prepared as described above and administered with dosing of 0.3 mg/kg or 3 mg/kg by i.p. injection three times a week; one cycle of treatment consisted of injections in the first and second week and no injection in the third week. Complete blood counts were determined by hemocytometer on day 26. Mice were euthanized after two cycles of treatment on day 33. Protein levels of MYC and GSPT1 were determined by flow cytometry or by western blot in bone marrow specimens.

### Cell lines, cell culture and generation of GT19630-resistant HL-60 cells

AML cell lines used in the study were obtained and cultured as previously described^1^. T-ALL and B-ALL cell lines were obtained and cultured as previously described^3,4^. DLBCL cell lines DLCL2, HT and Riva were kindly provided by Dr. Michael Green and were cultured in RPMI 1640 media supplemented with 10% FBS and with 100 mg/mL of penicillin and 100 U/mL of streptomycin at 37°C in 5% CO_2_ in a humidified atmosphere. All cell lines were subjected to cell line authentication performed by the Cytogenetics and Cell Authentication Core at The University of Texas MD Anderson Cancer Center and to mycoplasma testing every 6 months. HL-60 cells were subjected to chronic exposure to GT19630, with a concentration starting at 0.5 nM. The concentration of GT19630 was gradually increased to 10 nM by keeping their exponential growth and viabilities over 90% viability. Control HL-60 cells were treated with DMSO in the same manner as GT19630. The chronic exposure was performed with three biological replicates and resulted in GT19630-resistant (GTR) and -sensitive (GTS) HL-60 cells. All cell lines were submitted to DNA fingerprinting for authenticity.

### Reagents

Reagents used were G418 (Millipore-Sigma, A1720), CC-90009 (Selleckchem, S9832), ixazomib (MedChemExpress, HY-10452) bortezomib (MedChemExpress, HY-10227),

### Cancer cell line screening

A series of blood cancer, small cell lung cancer (SCLC), triple-negative breast cancer (TNBC) and glioma cell lines were treated with a series of concentrations of GT19630 and were subjected to a CellTiter-Glo assay by HD Biosciences.

### Development of MYC/GSPT1 degrader

GT19077, an inhibitor of MYC:MAX protein-protein interaction, was used as warhead. GT19406 was used as an E3 ligand. GT19077 and GT19406 were linked by flexible linkers to generate early MYC degrader GT19430. Cellular assay using HL-60 cells, where MYC is amplified^5^, for inhibiting proliferation (cell titer glo) and an ELISA for c-MYC protein levels were performed to investigate primary activities of candidate compounds. Then, cell proliferation inhibition and ELISA in TF-1 cells, where MYC is constitutively active by IL-5-JAK-STAT5 pathway^6^, were performed as counterscreening assays for candidate compounds (**Supplementary Fig. S3A**). During a structure–activity relationship study, the flexible linkers were replaced with rigid linkers to generate GT19506, an early lead molecule with improved potency and selectivity. GT19506 was further optimized with a novel CRBN E3 linker to improve potency, selectivity and in vivo potency and pharmacokinetic properties, leading to the generation of GT19630, the lead MYC protein degrader compound.

### Cell-free affinity pull-down assay

Recombinant MYC protein dilutions were aliquoted and stored on ice. Half of the were used as input. The remaining half were incubated with 5 µM of biotinylated GT19630 or different concentrations of naked GT19630 on ice for 2 hours. Next, 30 µL of Streptavidin Dynabeads (10 mg/mL) (Thermo Fisher Scientific, 11205D) were added, and were incubated on ice for 15–30 min. During magnet adsorption, samples were washed at least four times with 1 mL RIPA and PBS-T. SDS-PAGE was performed for the input and elute samples after boiling the samples at 100°C for 5 min. Western blotting was performed with MYC antibody (Abcam, ab32072).

### Plasmids and lentiviral transfection for MYC-stop-EGFP reporter assay

pLV[Exp]-Puro-EF1A>hMYC[NM_002467.6]*/P2A/EGFP plasmids with different stop codons (TGA or TAA) after the *MYC* gene were custom-designed and purchased from VectorBuilder. Each plasmid was transfected with packaging plasmids to produce lentivirus in HEK293T cells, and virus products were infected into HEK293T and HL-60 cells. Detailed lentivirus transfection was previously described^2^. Puromycin-selected HEK293T and HL-60 cells were treated with G418, CC-90009 or GT19630, and percentages of GFP-positive cells were determined by flow cytometry.

### Transfection of siRNAs

HL-60 cells were transfected with a pool of control siRNAs, *MYC* siRNAs or *GSPT1* siRNAs (Horizon Discovery; L-003282-02-0010 and LQ-019644-00-0002, respectively) by electroporation using a Nucleofector 2b device with Cell Line Nucleofector Kit L (Lonza) according to the manufacturer’s recommended protocol.

### Homogenous time-resolved fluorescence (HTRF) assay

HTRF assay was performed using HTRF Cereblon Binding Kits (Revvity, #64BDCRBNPEG) according to the manufacturer’s protocol. Briefly, the reference solution was transferred to a 384-reaction plate and compounds with a final concentration of 100 µM with a 3-fold concentration gradient were made. Human CRBN protein was transferred to the 384-reaction plate, and the plate was centrifuged at 1,000 rpm for 1 min and incubated at 25 °C for 15 min. Then, mixtures of either thalidomide red or GT19630 and Glutathione S-transferase conjugated with Europium cryptate was introduced into the 384-reaction plate, and the plate was centrifuged at 1,000 rpm for 1 min and incubated at 25 °C for 180 min. HTRF signals (ratio of 665/615 nm) were measured using a PHERAstar FSX microplate reader (BMG Labtech).

### Generation of CRBN knockout cells with inducible CRISPR-Cas9 system

The inducible cas9 plasmid (pCW-Cas9), a gift from Eric Lander & David Sabatini ^7^ was transfected into HEK293T cells along with the packaging vectors (pCMV + psPAX2) for the production of lentivirus particles. MOLM13 and HEK293T cells were transduced with cas9 lentivirus as per the already standardized protocol ^2^. Successfully transduced cells were selected using 1 ug/ml puromycin until the viability of surviving cells reached > 95%. The cells were sorted into single cell suspensions in a 96-well format using a Cytoflex cell sorter (Beckman Coulter, Brea, CA). Successfully grown clones were expanded and cas9 expression was induced using doxycycline (1 ug/ml for 48hrs followed by 24hr chase). The cas9 expression was verified using western blot using antibodies against cas9 (#14697, CST) β-actin (#4970, CST). Cas9 expressing MOLM13 and HEK293T clones without any leakage was selected for further studies. To generate knockout, pLKO5.sgRNA.EFS.GFP ^8^, kindly provided by Dr. Benjamin Ebert was used. Multiple sgRNA for CRBN were designed using CHOP-CHOP ^9^. Primers were designed by adding BsmBI-v2 overhangs, annealed, and treated with T4 PNK for 5’ end phosphorylation. Primer sequences are described below. The pLKO5.sgRNA.EFS.GFP vector was digested using BsmBI-v2 for 30 minutes, followed by phosphatase treatment. The digested plasmid was gel purified and used for annealing the sgRNA fragment. The correct clone was verified using Sanger sequencing and used for lentivirus production as described above. The resultant sgRNA lentivirus particles were transduced into MOLM13-Cas9 cells, followed by doxycycline induction. Single GFP-positive cells were sorted into 96-well plates using Bigfoot Spectral Cell Sorter (ThermoFisher Scientific, Waltham, MA). Successfully grown clones were expanded and CRBN knockout was confirmed by western blot.

Primer sequences

hCRBN-sgRNA-f2: CACCGTTACATACTGTATGTGATGT

hCRBN-sgRNA-r2: AAACACATCACATACAGTATGTAAC

hCRBN-sgRNA-f3: CACCGTTCTAATTGAACTGCAGACA

hCRBN-sgRNA-r3: AAACTGTCTGCAGTTCAATTAGAAC

### Seahorse analyses

Seahorse analyses were performed as previously described^10,11^. HL-60 GTS and GTR cells were treated with DMSO or 5 nM GT19630 for 12 hours, and cells were evaluated using an XF Analyzer (Agilent).

### Metabolomic and metabolic flux analyses

HL-60 GTS and GTR cells were treated with DMSO or 5 nM GT19630 for 12 hours in glucose-free and glutamine-free RPMI-1640 media (MilliporeSigma) supplemented with dialyzed FBS (MilliporeSigma) and with non-labeled glucose and glutamine, ^13^C_6_-labeled glucose and non-labeled glutamine, or non-labeled glucose and ^13^C_5_,^15^N_2_-labeled glutamine (Cambridge Isotope Laboratories). Cells were collected from each sample after washing with PBS twice. Then, 500 µL of supernatant was collected from each sample and filtered using a Nanosep 3K ultracentrifugal device (Pall Corp.) for 20 min at 4°C as previously descrbed^12^. Cells and supernatants were subjected to snap-freezing by liquid nitrogen for further mass spectrometry– based metabolomic analyses. Intracellular polar metabolites were extracted using a modified Bligh-Dyer method as previously reported^10,13,14^ and then analyzed using an ultra-high-pressure liquid chromatography (UHPLC) Vanquish tandem ultra-high-resolution Orbitrap IQ-X Tribrid mass spectrometer (Thermo Fisher Scientific). The mass spectrometer was calibrated for mass accuracy prior to analysis and continuously monitored during data acquisition to ensure mass accuracy remained below 5 ppm. A pooled quality control sample was injected after every six samples to assess instrument performance and ensure consistency across runs. The polar fraction was first analyzed using an Atlantis Premier BEH Z-HILIC 1.7-µm, 2.1 x 150 mm Column (Waters), and then re-analyzed using a Kinetex 2.6-µm C18, 150 x 2.1 mm Column (Phenomenex) as previously described^15^. The raw MS data were processed for peak picking and spectral alignment using the Sieve 2.2 application (Thermo Fisher Scientific). Peaks were then normalized using probabilistic quotient normalization. Metabolite identification was conducted using an in-house Matlab script to match accurate masses and retention times to an in-house standards library, including the IROA MS metabolite library (IROA Technologies). Accurate masses were also matched against the Human Metabolome Database and the Kyoto Encyclopedia of Genes and Genomes. Isotope natural abundance correction was performed using the AccuCor2 tool^16^. Statistical significance was calculated using two-tailed Student *t* test.

### Single-cell mass cytometry (CyTOF)

Frozen primary AML and normal bone marrow samples from healthy donors were thawed and incubated as previously described^1^. Washed cells were co-cultured with mesenchymal stromal cells and treated with DMSO or 64 nM GT19630 for 24 hours, followed by staining procedures with metal-tagged antibodies as previously described^1^. Antibodies used in the present study were CD45 (89 Yb, clone HI30, Standard BioTools, #3089003B), CD4 (151 Eu, clone A161A1, BioLegend, #357402), CD8 (162 Dy, clone RPA-T8, Standard BioTools, #3162015B), CD11b (148 Nd, clone M1/70, BioLegend, #3148003B), CD33 (158 Gd, clone WM53, Standard BioTools, #3158001B), CD34 (164 Dy, clone 581, BioLegend, #343502), CD38 (167 Er, HIT2, BioLegend, #303502), c-MYC (169 Tm, clone Y69, Abcam, #ab168727), Ki-67 (168 Er, clone BLR021E, Bethyl Laboratories, #A700-021CF), CD19 (161 Dy, clone HIB19, BioLegend, #302202), CD44 (166 Er, clone BJ18, Standard BioTools, #3166001B), CD56 (163 Dy, clone NCAM16.2, Standard BioTools, #3163007B), CD90 (171 Yb, clone 5E10, BioLegend, #328102), CD99 (165 Ho, clone TU12, BD Biosciences, #555687), CD123 (145 Nd, clone 7G3, BD Biosciences, #554527), CD45RA (170 Er, clone HI100, BioLegend, #304102), CD47 (141 Pr, clone CC2C6, BioLegend, #323102), CLL-1 (174 Yb, clone 50C1, BioLegend, #353602), cleaved PARP (143 Nd, clone F21-852, Standard BioTools, #3143011A), g-H2A.X (147 Sm, clone BLR053F, Bethyl Laboratories, #A700-021CF), BAX (173 Yb, clone 2D2, BioLegend, #633602), p-AKT (159 Tb, clone M89-61, Standard BioTools #560397), p-STAT5 (149 Sm, clone C71E5, Cell Signaling Technology, #9314BF), p-STAT3 (152 Sm, clone 4/P-STAT3, BD Biosciences, #612357), MCL-1 (176 Yb, D2W9E, Cell Signaling Technology, #94296BF), p21 (153 Eu, clone 12D1, Cell Signaling Technology, #2947BF), Bcl-2 (144 Nd, clone 100, BioLegend, #658702), ERK1 (139 La, clone 250603, R&D Systems, #MAB1940), cyclin D1 (142 Nd, clone G124-326, BD Biosciences, #554180), CXCR4 (175 Lu, clone 12G5, Standard BioTools, #3175001B) and PD-L1 (172 Yb, clone 29E.2A3, BioLegend, #329702).

### Apoptosis analyses

Apoptosis analyses using annexin V (APC, BioLegend, #640941) and DAPI staining assays using counting beads (AccuCount, Spherotech, #ACFP-70-10) to determine absolute live cell numbers. For primary AML samples, we used surface antibodies for human CD45 (PE, BD Biosciences, #560975), CD34 (PE-Cy7, BioLegend, #343516), CD38 (APC Fire 750, BioLegend, #356626), CD117 (PE Dazzle 594, BioLegend, #313226), CD123 (APC, BioLegend, #306012), annexin V (PerCP Cy5.5, BioLegend, #640936) and DAPI. Compusyn software was used to calculate IC50 values.

### Immunoblot analyses

Immunoblot analyses were performed as previously described^1^. Antibodies used in the present study were MYC (clone Y69, Abcam, #ab32072), GSPT1 (clone EPR22908-103, Abcam #ab234433), ATF-3 (clone D2Y5W, Cell Signaling Technology, #33593), ATF-4 (clone D4B8, Cell Signaling Technology, #11815), MCL-1 (D35A5, Cell Signaling Technology, #5453), α-tubulin (clone 11H10, Cell Signaling Technology, #2125), GAPDH (clone 10B13, MilliporeSigma, #ZRB374), histone H3 (Cell Signaling Technology, #9715) and β-actin (MilliporeSigma, #A5316).

### Quantitative RT-PCR

The mRNA expression levels in HL-60 cells transfected with siRNAs for *MYC* and *GSPT1* were quantified using TaqMan gene expression assays, including MYC (Hs00153408_m1), MCM4 (Hs00907398_m1) and B2M (Hs00187842_m1). B2M was used as the internal control.

### RNA sequencing

HL-60 GTS cells were treated with DMSO or 5 nM GT19630 for 12 hours in three biological replicates and total RNA was extracted from each sample using an RNeasy extraction kit (QIAGEN, #74104) following the manufacturer’s protocol. The RNA-sequencing (RNA-seq) analysis pipeline was performed as previously described^17^.

### Differential gene expression profiles and pathway analyses

Differential gene expression profiling was performed as previously described^1^. Gene set enrichment analysis (GSEA) was performed using GSEA v4.3.3 software from the Broad Institute with Molecular Signatures Database gene sets (v2024.1)^18^ to compare differential gene expression profiles of HL-60 GTS cells treated with DMSO or 5 nM GT19630 for 12 hours.

### DNA sequencing

DNA samples from HL-60 and Raji cells were extracted using a QIAamp DNA Blood Mini Kit (QIAGEN, #51104). We first amplified the *MYC* gene by PCR the forward primer 5’ TACTGCGACGAGGAGGAGAA-3’ and the reverse primer 5’-AGAGGGTAGGGGAAGACCAC-3’, and amplified DNA samples were subjected to direct sequencing using the forward primer. ApE software was used to analyze the sequencing data^19^.

### Quantification and statistical analyses

Analyses using the Munich Leukemia Laboratory dataset were performed as previously reported ^20^. In the AML dataset, MYC expression levels were classified into four groups (Quadrant (Q) 1, highest; Q2, second highest; Q3, second lowest; and Q4, lowest), and multivariate analyses were used to detect statistically significant differences between proportions of Q1 and Q4 according to genetic or cytogenetic variations. The processed data in GSM935516, GSM935643, GSM822290, GSM822291, GSM1006866, and GSM1003607 were downloaded from Gene Expression Omnibus and MYC peaks for *GSPT1* and *ETF1* genes were visualized using IGV genome browser. Dependency scores in cancer cell lines determined by CRISPR screening were obtained from the gene essentiality viewer using shinydepmap^22^. The two-tailed Student *t*-test was used to compare two groups unless otherwise specified. One-way analysis of variance (ANOVA) and the Bonferroni ad hoc test were used to compare more than two groups. The Kaplan-Meier method was used to generate survival curves, and the log-rank test was used to compare the survival times of two groups. A *P* value of less than 0.05 was considered statistically significant. All experiments were performed with three or more biological replicates, and error bars in graphs represent mean ± standard deviation (SD) unless otherwise specified.

## Supplementary figures and legends

**Supplementary Fig. S1.**
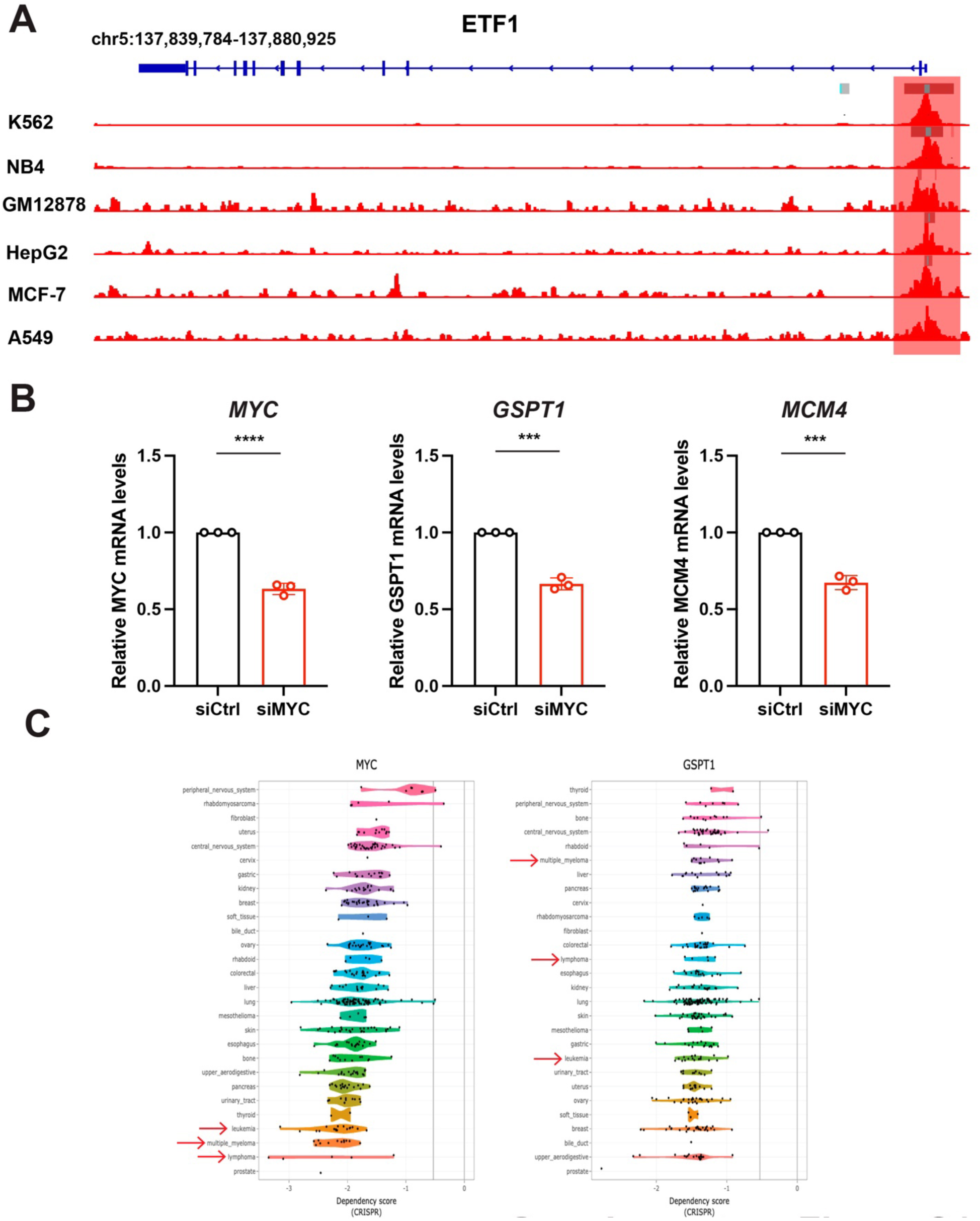
MYC regulates genes involved in translation termination and MYC and GSPT1 are essential in tumor cells. **A.** Binding of MYC to the TSS of ETF1 in hematologic and solid tumor cells. ETF1 gene map at chromosome 5 is shown on top. Significantly enriched regions are highlighted with gray bars above peaks. Highest peaks are highlighted in light blue. **B.** Relative quantitation of *MYC*, *GSPT1* and *MCM4* after introducing scramble control siRNA (siCtrl) and siRNA for MYC (siMYC) in HL-60 cells. Beta-2-microglobulin was used as an internal control. *** P < 0.001, **** P < 0.0001. **C.** Dependency scores, where lower scores represent bigger lethality when the subject gene is knocked our for cancer cell lines, for MYC and GSPT1 determined by CRISPR screening among cancer cell lines from DepMap datasets. Cell lines representing leukemia, lymphoma and multiple myeloma are annotated with red arrows.

**Supplementary Fig. S2.**
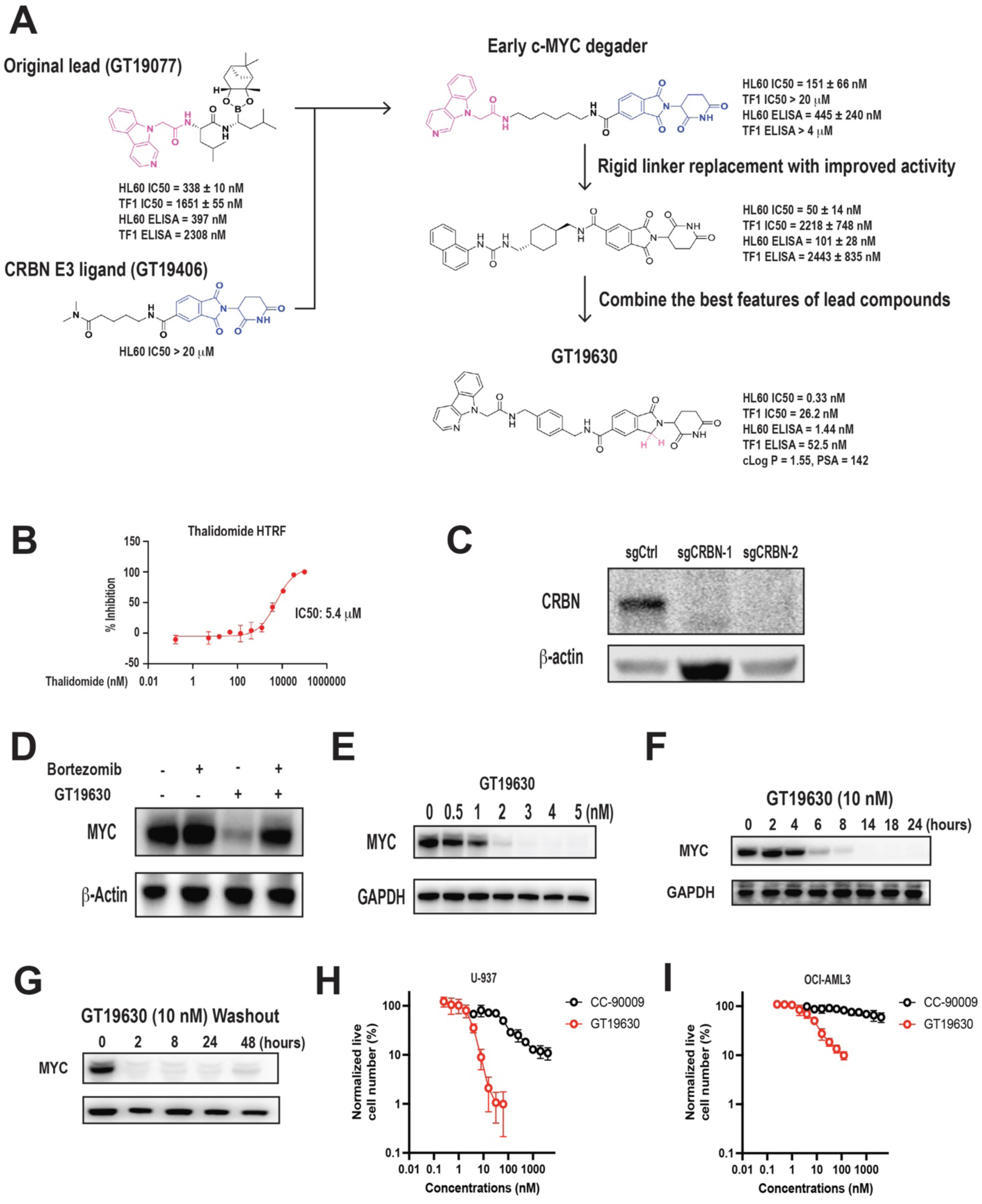
Development of MYC/GSPT1 PROTAC and basic characteristics of GT19630. **A.** Schematic diagram of the development of GT19630. PSA, polar surface area. **B.** IC50 values for CRBN binding of thalidomide and GT19630 as determined by homogenous time-resolved fluorescence assay. **C.** Homogeneous time-resolved fluorescence (HTRF) assays for the interaction of CRBN with thalidomide. **D.** Western blot of MYC after treatment with DMSO (shown as -), 80 nM bortezomib, 5 nM GT19630 and both bortezomib and GT19630 in HL-60 cells for 8 hours. β-actin served as a loading control. **E.** MYC protein levels in HL-60 cells treated with the indicated concentrations of GT19630. GAPDH served as a loading control. **F.** MYC protein levels in HL-60 cells treated with 10 nM GT19630 for the indicated time periods. GAPDH served as a loading control. **G.** MYC protein levels in HL-60 cells treated with 10 nM GT19630 for 24 hours and washed out. GAPDH served as a loading control. **H, I.** Normalized live cell numbers of U-937 cells (**H**) and OCI-AML3 cells (**I**) treated with indicated concentrations of GT19630 or CC-90009 for 48 hours.

**Supplementary Fig. S3.**
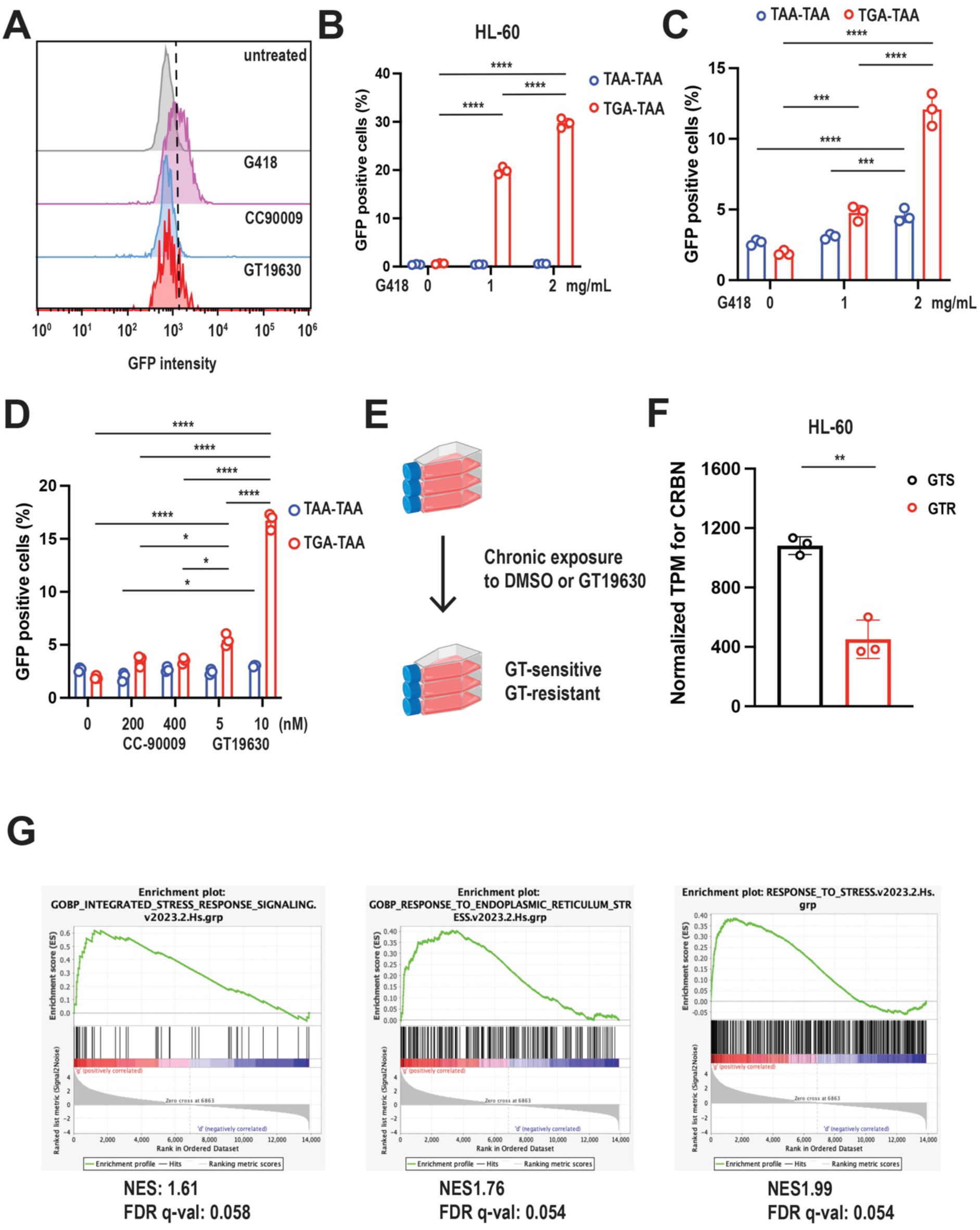
MYC/GSPT1 degradation induces stop codon readthrough and integrated stress response (ISR). **A.** Representative histograms of GFP intensities of live (annexin V/DAPI–negative) HL-60 cells treated with 2 mg/mL G418, 200 nM CC-90009 or 10 nM GT19630. **B.** Percentages of GFP-positive HL-60 cells transduced with two different MYC-stop-GFP constructs and treated with indicated concentrations of G418. **C.** Percentages of GFP-positive HL-60 cells transduced with two different MYC-stop-GFP constructs and treated with indicated concentrations of G418. **D.** Percentages of GFP-positive HEK293T cells after 24 hours of treatment with CC-90009, GT19630 or no drug (0-nM control). **E.** Schematic diagram of HL-60 cells undergoing chronic exposure to DMSO or GT19630 to obtain GT-sensitive and - resistant cells. **F.** Normalized transcripts of CRBN in HL-60 GT19630-sensitive (GTS) and - resistant (GTR) cells. **G.** Gene set enrichment analyses for the pathways Integrated Stress Response Signaling (left), Response to Endoplasmic Reticulum Stress (middle) and Response to Stress (right). NES, normalized enrichment score. \**P* < 0.05, ** P < 0.01, \*\*\**P* < 0.001, \*\*\*\**P* < 0.0001.

**Supplementary Fig. S4.**
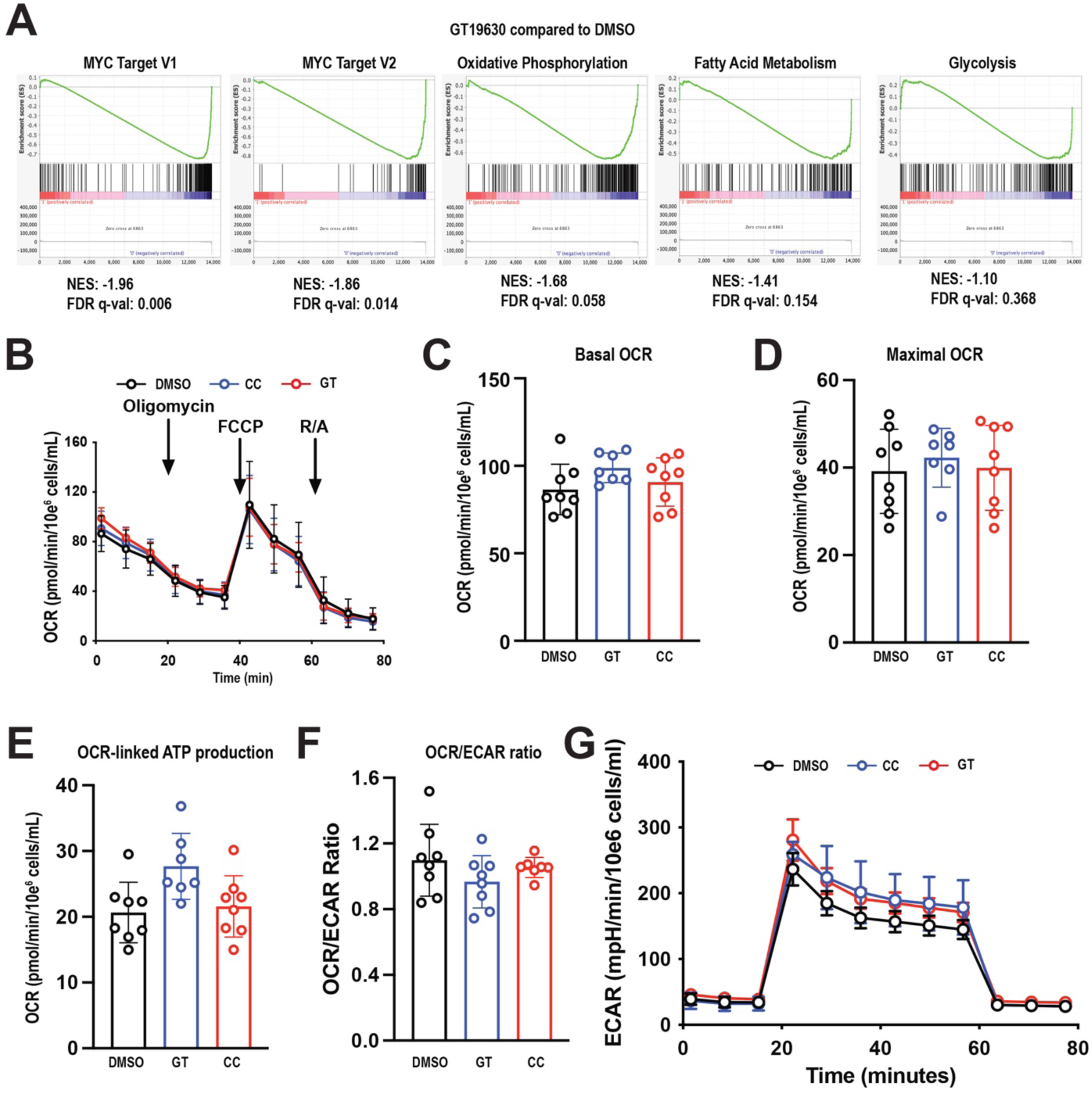
Impact of GT19630 on metabolic pathways. **A.** Gene set enrichment analyses for pathways for MYC Targets, Oxidative Phosphorylation, Fatty Acid Metabolism and Glycolysis in HL-60 cells treated with 5 nM GT19630 for 12 hours. FDR q-values < 0.25 is considered significant. **B.** Oxygen consumption rate (OCR) measured by Seahorse analysis in HL-60 GTR cells treated with DMSO, 100 nM CC-90009 (CC) or 5 nM GT19630 (GT) for 12 hours. **C-F.** Measures of basal OCR (**C**), maximal OCR (**D**), OCR-linked ATP production (**E**) and OCR/ECAR ratio (**F**) in HL-60 GTR cells treated with DMSO, 100 nM CC or 5 nM GT for 12 hours. **G.** Extracellular acidification rate (ECAR) measured by Seahorse analysis in HL-60 GTR cells treated with DMSO, 100 nM CC or 5 nM GT for 12 hours.

**Supplementary Fig. S5.**
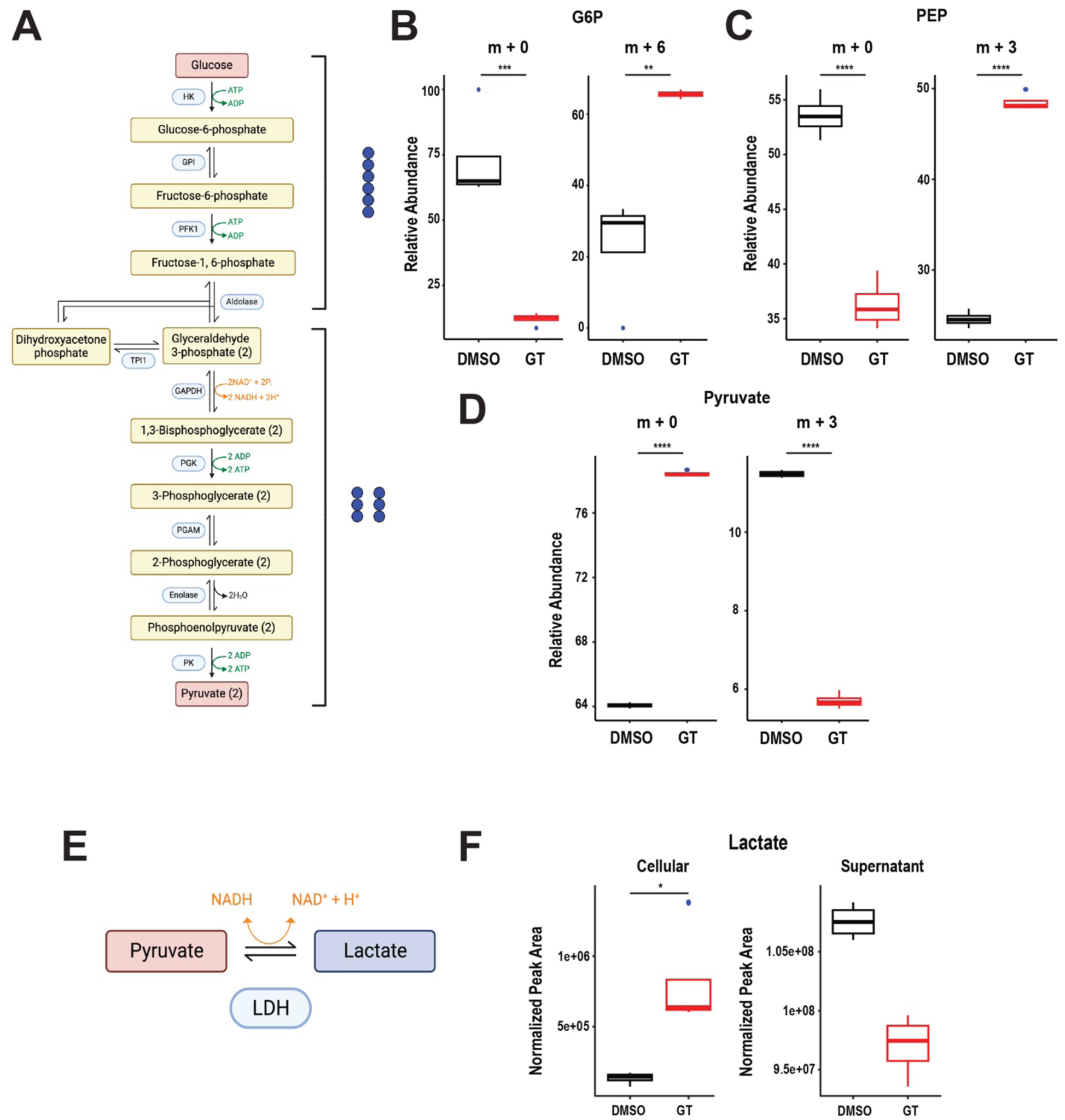
MYC/GSPT1 degradation accumulates glycolysis intermediates and cellular lactate. **A.** Map of the TCA cycle for metabolic flux analyses (MFA). ^13^C_6_-labeled carbon is shown as blue circles at the right. **B-D, F.** Levels of intermediates in HL-60 GTS cells treated with DMSO or 5 nM GT19630 (GT) for 12 hours. **B.** m+0 and m+6 glucose-6-phosphate (G6P) levels. **C.** m+0 and m+3 phosphoenolpyruvate (PEP) levels. **D.** m+0 and m+3 pyruvate levels. **E.** Schematic diagram of the pyruvate and lactate reaction for MFA. **F.** Cellular and supernatant lactate levels. \**P* < 0.05, \*\**P* < 0.01, \*\*\**P* < 0.001, \*\*\*\**P* < 0.0001.

**Supplementary Fig. S6.**
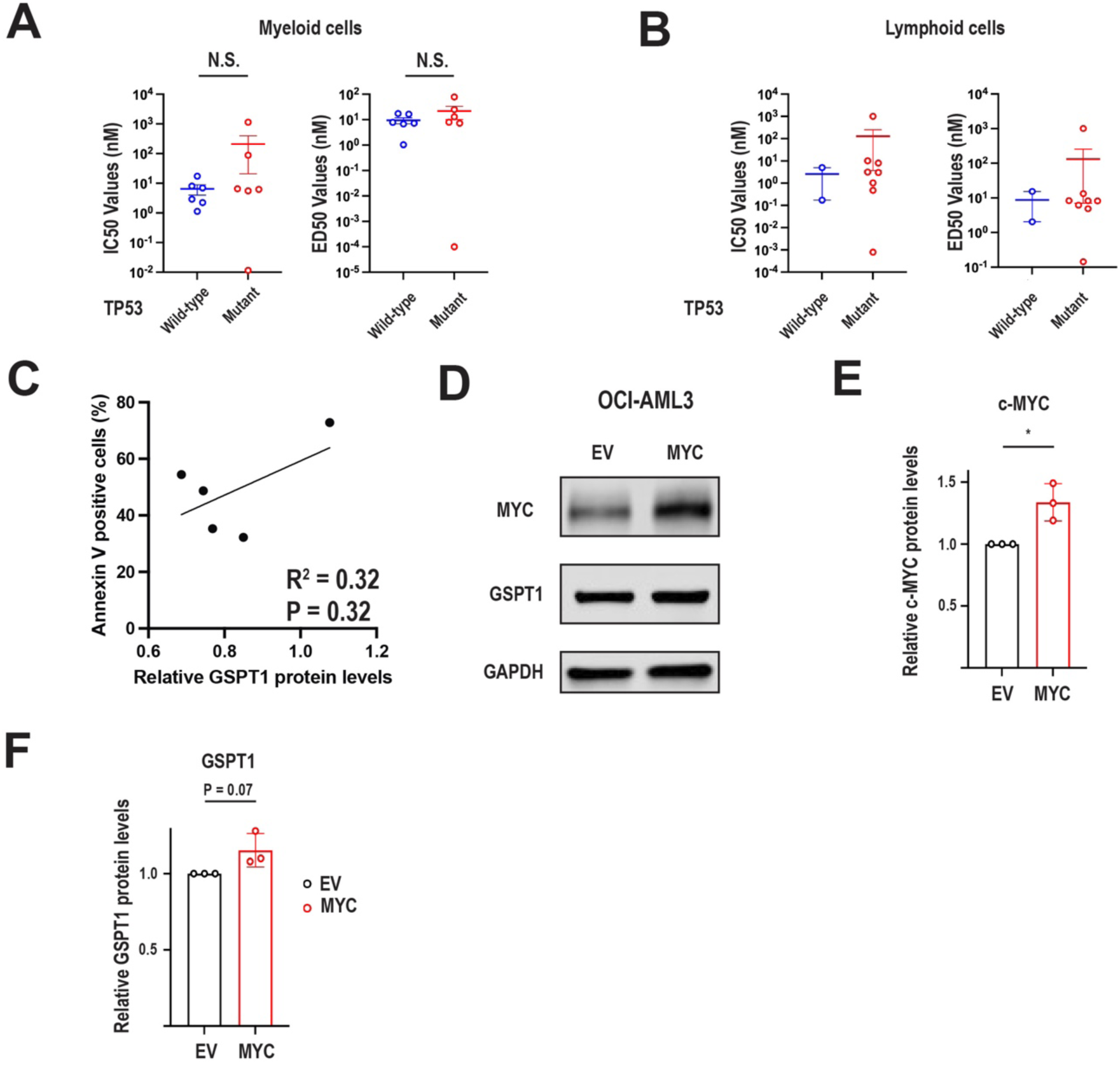
GT19630 induced TP53 independent cell death in blood cancer cells. **A.** IC50 and ED50 values of *TP53* wild-type and mutant myeloid tumor cell lines. **B.** IC50 and ED50 values of *TP53* wild-type and mutant lymphoid tumor cell lines. **C.** Correlation plot of baseline protein levels of GSPT1 and percentages of annexin V–positive cells in MOLM-13 *TP53* knockout and mutant cells treated with 40 nM GT19630 for 48 hours. **D.** Protein levels of MYC and GSPT1 in OCI-AML3 cells transduced with empty vector (EV) or MYC-overexpressing (MYC) constructs. GAPDH served as a loading control. **E.** Protein levels of c-MYC in OCI-AML3 EV and MYC cells. **F.** Protein levels of GSPT1 in OCI-AML3 EV and MYC cells. N.S., not significant. \**P* < 0.05.

**Supplementary Fig. S7.**
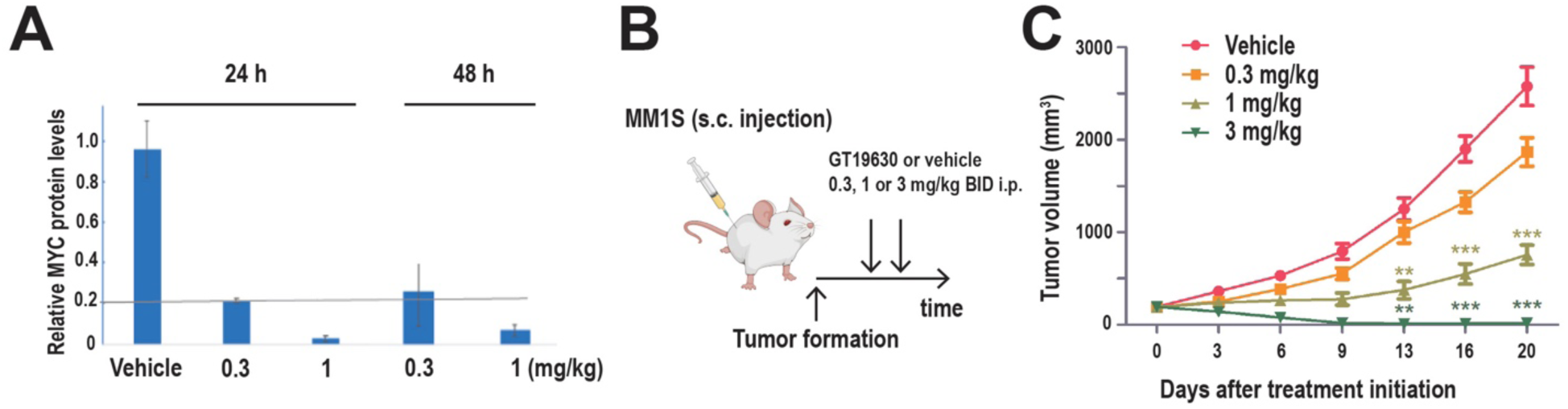
**A.** Relative protein levels of MYC in HL-60 tumor samples (N = 3 for each group) obtained from mice 24 and 48 hours after treatment with vehicle or indicated doses of GT19630. **B.** Schematic diagram of in vivo experiment. Mice were subcutaneously (s.c.) injected with MM1S cells and treated with indicated doses of GT19630 or vehicle once a day. **C.** Size of MM1S tumors in each treatment group at indicated days.

**Supplementary Fig. S8.**
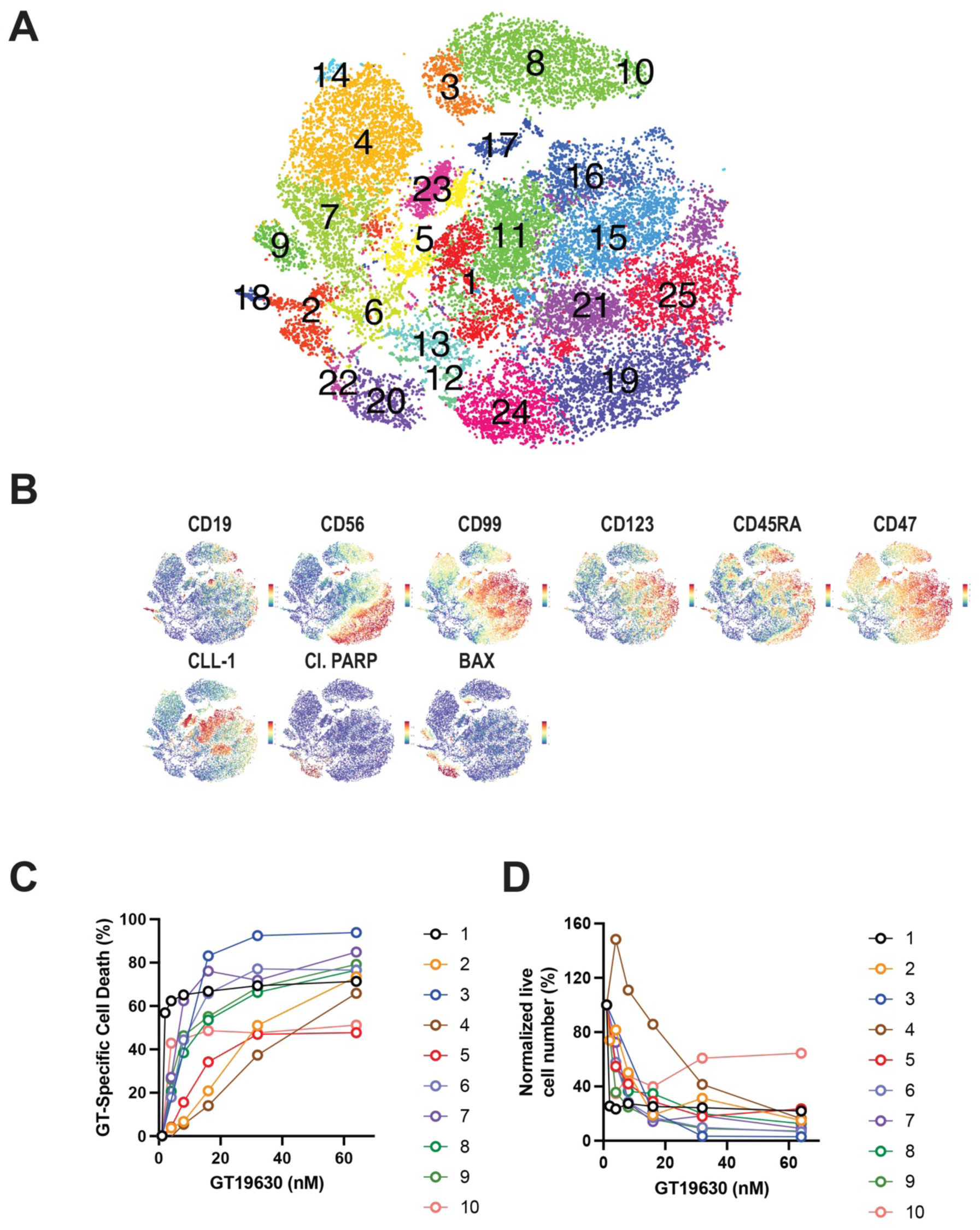
Single-cell mass cytometry (CyTOF) in normal bone marrow and primary AML cells and efficacy of GT19630 in primary AML samples. **A.** Unsupervised clustering of two normal bone marrow and three AML samples subjected to single-cell mass cytometry. **B.** Indicated protein levels determined by CyTOF in clusters after unsupervised clustering. **C.** Percentages of annexin V– and/or DAPI-positive cells of 10 primary AML samples treated with indicated concentrations of GT19630 for 72 hours. **D.** Normalized numbers of live cells for the 10 primary AML samples after GT19630 treatment for 72 hours.

**Supplementary Fig. S9.**
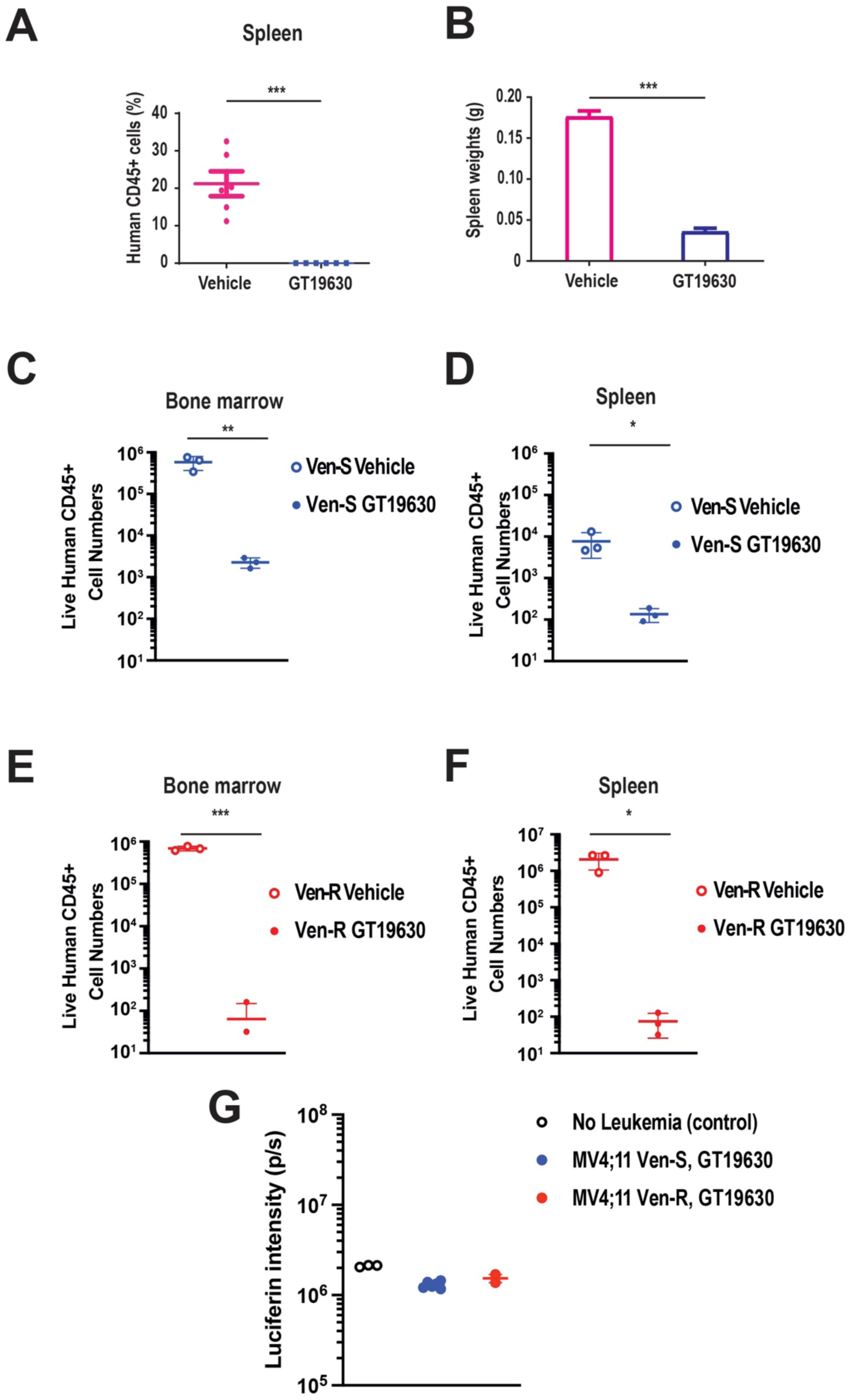
In vivo activities of GT19630 in AML models. **A.** Percentages of human CD45-positive cells in spleens of NOD/SCID mice injected with AML PDX cells and treated with vehicle or GT19630. **B.** Weights of spleens of mice on day 15 of treatment with vehicle or GT19630. **C-F.** Live human CD45-positive cell numbers in bone marrow or spleen samples from NSG mice injected with MV4;11 Ven-S or Ven-R cells and treated with vehicle or GT19630. **C.** Bone marrow samples; Ven-S cells. **D.** Spleen samples; Ven-S cells. **E.** Bone marrow samples; Ven-R cells. **F.** Spleen samples; Ven-R cells. **G.** Luciferin intensities determined by bioluminescence imaging at day 175 in NSG mice without AML cells (No Leukemia (control)) or NSG mice injected with MV4;11 Ven-S cells or Ven-R cells and treated with GT19630. \**P* < 0.05, \*\**P* < 0.01, \*\*\**P* < 0.001.

**Supplementary Fig. S10.**
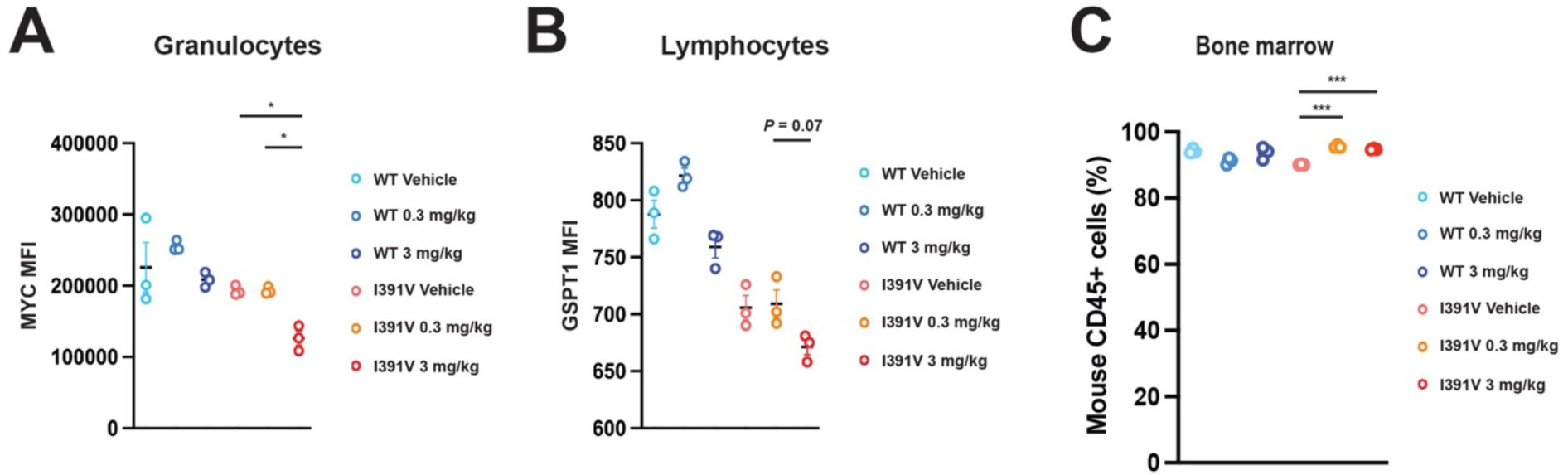
GT19630 reduces MYC and GSPT1 levels in humanized *Crbn^I391V^*mice. **A, B.** Protein levels of MYC in granulocytes (**A**) and of GSPT1 in lymphocytes (**B**) determined by flow cytometry in bone marrow samples of CL57B/6 mice with wild-type (WT) and I391V mutant *Crbn* that received the indicated in vivo treatments. **C.** Percentages of CD45-positive cells in bone marrow samples of CL57B/6 mice with wild-type (WT) and I391V mutant *Crbn* that received the indicated in vivo treatments. All samples were collected on day 33 after treatment initiation.

**Supplementary Table S1.**
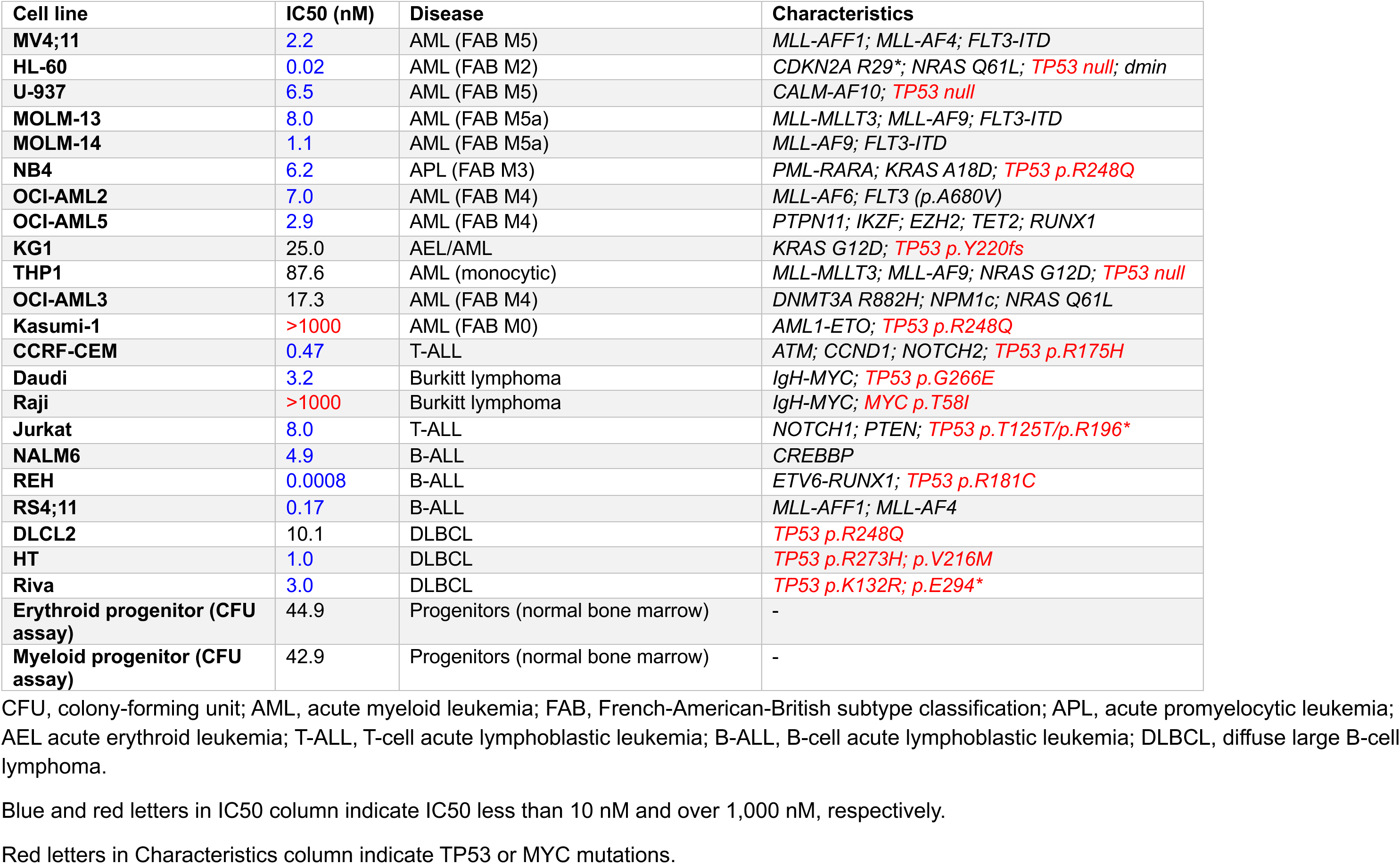
IC50 values and characteristics of leukemia and lymphoma cell lines treated with GT19630.

**Supplementary Table S2.** Characteristics of primary AML samples used for GT19630 treatment.

| ID | Age (yr) | Sex | Status | # of Tx | Blasts | ELN criteria | Mutations | Cytogenetics |
| --- | --- | --- | --- | --- | --- | --- | --- | --- |
| 1 | 43 | F | Ref | 6 | 89% | Adverse | *BCORL1* | Complex with t(9;11)(p22;q23) |
| 2 | 57 | F | Ref | 2 | 82% | Adverse | *TP53* (p.S215T/p.I195T) | Complex |
| 3 | 81 | M | Rel | 1 | 86% | Adverse | *BCOR*, *CBL*, *DNMT3A*, *TET2*, *TP53* | 45,X,-Y[3]/46,XY[17] |
| 4 | 68 | F | Rel | 2 | 95% | Adverse | *TP53* (p.R175H) | Complex, extra copy of MYC |
| 5 | 63 | F | Ref | 2 | 24% | Adverse | *TP53* (p.L130fs) | Complex |
| 6 | 77 | M | Ref | 3 | 49% | Intermediate | *JAK2*, *RUNX1*, *TET2* | 48,XY,+8,+21[20] |
| 7 | 69 | F | Ref | 4 | 88% | Adverse | *KRAS*, *RUNX1* | Complex with t(3;21)(q26.2;q22) |
| 8 | 71 | F | Ref* | 3 | 59% | N.A. | *ASXL2*, *NRAS*, *SRSF2* | Complex with inv(3)(q21q26.2) |
| 9 | 71 | M | Ref** | 2 | 81% | Intermediate | N.A. | 44,idem,+mar2,2dmin[1] |
| 10 | 78 | F | Rel | 1 | 70% | Adverse | *JAK2*, *TP53* (p.C124*), *U2AF1* | 46,XX,-12,+mar[1]/46,XX[9] |

Ref, refractory; Rel, relapsed; Tx, treatments; ELN, European LeukemiaNet genetic risk classification; N.A., not assessed.
\*This patient was diagnosed with myeloproliferative neoplasm, and the disease transformed to mixed-phenotype acute leukemia.
\*\*This patient was diagnosed with acute lymphoblastic leukemia and then developed myelodysplastic syndrome. The disease transformed to acute myeloid leukemia.

## References

1 Dang, C. V. et al. The c-Myc target gene network. Semin Cancer Biol 16, 253–264 (2006). 10.1016/j.semcancer.2006.07.014

2 Llombart, V. & Mansour, M. R. Therapeutic targeting of "undruggable" MYC. EBioMedicine 75, 103756 (2022). 10.1016/j.ebiom.2021.103756

3 Dhanasekaran, R. et al. The MYC oncogene - the grand orchestrator of cancer growth and immune evasion. Nat Rev Clin Oncol 19, 23–36 (2022). 10.1038/s41571-021-00549-2

4 Manolov, G. & Manolova, Y. Marker band in one chromosome 14 from Burkitt lymphomas. Nature 237, 33–34 (1972). 10.1038/237033a0

5 Dalla-Favera, R. et al. Human c-myc onc gene is located on the region of chromosome 8 that is translocated in Burkitt lymphoma cells. Proc Natl Acad Sci U S A 79, 7824–7827 (1982). 10.1073/pnas.79.24.7824

6 Campo, E. MYC in DLBCL: partners matter. Blood 126, 2439–2440 (2015). 10.1182/blood-2015-10-671362

7 Sanchez-Martin, M. & Ferrando, A. The NOTCH1-MYC highway toward T-cell acute lymphoblastic leukemia. Blood 129, 1124–1133 (2017). 10.1182/blood-2016-09-692582

8 Felsher, D. W. MYC Inactivation Elicits Oncogene Addiction through Both Tumor Cell-Intrinsic and Host-Dependent Mechanisms. Genes Cancer 1, 597–604 (2010). 10.1177/1947601910377798

9 van Galen, P. et al. Single-Cell RNA-Seq Reveals AML Hierarchies Relevant to Disease Progression and Immunity. Cell 176, 1265–1281 e1224 (2019). 10.1016/j.cell.2019.01.031

10 Nishida, Y. et al. Enhanced TP53 reactivation disrupts MYC transcriptional program and overcomes venetoclax resistance in acute myeloid leukemias. Sci Adv 9, eadh1436 (2023). 10.1126/sciadv.adh1436

11 Bekes, M., Langley, D. R. & Crews, C. M. PROTAC targeted protein degraders: the past is prologue. Nat Rev Drug Discov 21, 181–200 (2022). 10.1038/s41573-021-00371-6

12 Sasso, J. M. et al. Molecular Glues: The Adhesive Connecting Targeted Protein Degradation to the Clinic. Biochemistry 62, 601–623 (2023). 10.1021/acs.biochem.2c00245

13 Kozicka, Z. & Thoma, N. H. Haven’t got a glue: Protein surface variation for the design of molecular glue degraders. Cell Chem Biol 28, 1032–1047 (2021). 10.1016/j.chembiol.2021.04.009

14 Embree, C. M., Abu-Alhasan, R. & Singh, G. Features and factors that dictate if terminating ribosomes cause or counteract nonsense-mediated mRNA decay. J Biol Chem 298, 102592 (2022). 10.1016/j.jbc.2022.102592

15 Matyskiela, M. E. et al. A novel cereblon modulator recruits GSPT1 to the CRL4(CRBN) ubiquitin ligase. Nature 535, 252–257 (2016). 10.1038/nature18611

16 Sellar, R. S. et al. Degradation of GSPT1 causes TP53-independent cell death in leukemia while sparing normal hematopoietic stem cells. J Clin Invest 132 (2022). 10.1172/JCI153514

17 Surka, C. et al. CC-90009, a novel cereblon E3 ligase modulator, targets acute myeloid leukemia blasts and leukemia stem cells. Blood 137, 661–677 (2021). 10.1182/blood.2020008676

18 Chang, Y. et al. The orally bioavailable GSPT1/2 degrader SJ6986 exhibits in vivo efficacy in acute lymphoblastic leukemia. Blood 142, 629–642 (2023). 10.1182/blood.2022017813

19 Stine, Z. E., Walton, Z. E., Altman, B. J., Hsieh, A. L. & Dang, C. V. MYC, Metabolism, and Cancer. Cancer Discov 5, 1024–1039 (2015). 10.1158/2159-8290.CD-15-0507

20 Dong, Y., Tu, R., Liu, H. & Qing, G. Regulation of cancer cell metabolism: oncogenic MYC in the driver’s seat. Signal Transduct Target Ther 5, 124 (2020). 10.1038/s41392-020-00235-2

21 Bae, S. et al. MYC-mediated early glycolysis negatively regulates proinflammatory responses by controlling IRF4 in inflammatory macrophages. Cell Rep 35, 109264 (2021). 10.1016/j.celrep.2021.109264

22 Lee, K. M. et al. MYC and MCL1 Cooperatively Promote Chemotherapy-Resistant Breast Cancer Stem Cells via Regulation of Mitochondrial Oxidative Phosphorylation. Cell Metab 26, 633–647 e637 (2017). 10.1016/j.cmet.2017.09.009

23 Huh, Y. O. et al. Double minute chromosomes in acute myeloid leukemia, myelodysplastic syndromes, and chronic myelomonocytic leukemia are associated with micronuclei, MYC or MLL amplification, and complex karyotype. Cancer Genet 209, 313–320 (2016). 10.1016/j.cancergen.2016.05.072

24 Shimada, K., Bachman, J. A., Muhlich, J. L. & Mitchison, T. J. shinyDepMap, a tool to identify targetable cancer genes and their functional connections from Cancer Dependency Map data. Elife 10 (2021). 10.7554/eLife.57116

25 Baradaran-Heravi, A. et al. Effect of small molecule eRF3 degraders on premature termination codon readthrough. Nucleic Acids Res 49, 3692–3708 (2021). 10.1093/nar/gkab194

26 Wangen, J. R. & Green, R. Stop codon context influences genome-wide stimulation of termination codon readthrough by aminoglycosides. Elife 9 (2020). 10.7554/eLife.52611

27 Haertle, L. et al. Cereblon enhancer methylation and IMiD resistance in multiple myeloma. Blood 138, 1721–1726 (2021). 10.1182/blood.2020010452

28 Chen, Y. J. et al. Lactate metabolism is associated with mammalian mitochondria. Nat Chem Biol 12, 937–943 (2016). 10.1038/nchembio.2172

29 de la Cruz-Lopez, K. G., Castro-Munoz, L. J., Reyes-Hernandez, D. O., Garcia-Carranca, A. & Manzo-Merino, J. Lactate in the Regulation of Tumor Microenvironment and Therapeutic Approaches. Front Oncol 9, 1143 (2019). 10.3389/fonc.2019.01143

30 Cao, X. et al. Analysis of 3760 hematologic malignancies reveals rare transcriptomic aberrations of driver genes. Genome Med 16, 70 (2024). 10.1186/s13073-024-01331-6

31 Bhatia, K. et al. Point mutations in the c-Myc transactivation domain are common in Burkitt’s lymphoma and mouse plasmacytomas. Nat Genet 5, 56–61 (1993). 10.1038/ng0993-56

32 Kalkat, M. et al. MYC Deregulation in Primary Human Cancers. Genes (Basel) 8 (2017). 10.3390/genes8060151

33 Boettcher, S. et al. A dominant-negative effect drives selection of TP53 missense mutations in myeloid malignancies. Science 365, 599–604 (2019). 10.1126/science.aax3649

34 Bulaeva, E. et al. MYC-induced human acute myeloid leukemia requires a continuing IL-3/GM-CSF costimulus. Blood 136, 2764–2773 (2020). 10.1182/blood.2020006374

35 Ohanian, M. et al. MYC protein expression is an important prognostic factor in acute myeloid leukemia. Leuk Lymphoma 60, 37–48 (2019). 10.1080/10428194.2018.1464158

36 Yun, S. et al. Prognostic significance of MYC oncoprotein expression on survival outcome in patients with acute myeloid leukemia with myelodysplasia related changes (AML-MRC). Leuk Res 84, 106194 (2019). 10.1016/j.leukres.2019.106194

37 Sachdeva, M. et al. p53 represses c-Myc through induction of the tumor suppressor miR-145. Proc Natl Acad Sci U S A 106, 3207–3212 (2009). 10.1073/pnas.0808042106

38 Porter, J. R. et al. Global Inhibition with Specific Activation: How p53 and MYC Redistribute the Transcriptome in the DNA Double-Strand Break Response. Mol Cell 67, 1013–1025 e1019 (2017). 10.1016/j.molcel.2017.07.028

39 Muller-Tidow, C. et al. Translocation products in acute myeloid leukemia activate the Wnt signaling pathway in hematopoietic cells. Mol Cell Biol 24, 2890–2904 (2004). 10.1128/MCB.24.7.2890-2904.2004

40 Tashiro, H. et al. Treatment of Acute Myeloid Leukemia with T Cells Expressing Chimeric Antigen Receptors Directed to C-type Lectin-like Molecule 1. Mol Ther 25, 2202–2213 (2017). 10.1016/j.ymthe.2017.05.024

41 Pei, S. et al. Monocytic Subclones Confer Resistance to Venetoclax-Based Therapy in Patients with Acute Myeloid Leukemia. Cancer Discov 10, 536–551 (2020). 10.1158/2159-8290.CD-19-0710

42 Kronke, J. et al. Lenalidomide induces ubiquitination and degradation of CK1alpha in del(5q) MDS. Nature 523, 183–188 (2015). 10.1038/nature14610

43 Lourenco, C. et al. MYC protein interactors in gene transcription and cancer. Nat Rev Cancer 21, 579–591 (2021). 10.1038/s41568-021-00367-9

44 Ruggero, D. The role of Myc-induced protein synthesis in cancer. Cancer Res 69, 8839–8843 (2009). 10.1158/0008-5472.CAN-09-1970

45 Pelletier, J., Graff, J., Ruggero, D. & Sonenberg, N. Targeting the eIF4F translation initiation complex: a critical nexus for cancer development. Cancer Res 75, 250–263 (2015). 10.1158/0008-5472.CAN-14-2789

46 Nishida, Y. et al. Inhibition of translation initiation factor eIF4a inactivates heat shock factor 1 (HSF1) and exerts anti-leukemia activity in AML. Leukemia 35, 2469–2481 (2021). 10.1038/s41375-021-01308-z

47 Hoelzer, D. et al. Improved outcome of adult Burkitt lymphoma/leukemia with rituximab and chemotherapy: report of a large prospective multicenter trial. Blood 124, 3870–3879 (2014). 10.1182/blood-2014-03-563627

48 Evens, A. M. et al. Burkitt lymphoma in the modern era: real-world outcomes and prognostication across 30 US cancer centers. Blood 137, 374–386 (2021). 10.1182/blood.2020006926

49 Abdallah, N. et al. Implications of MYC Rearrangements in Newly Diagnosed Multiple Myeloma. Clin Cancer Res 26, 6581–6588 (2020). 10.1158/1078-0432.CCR-20-2283

50 Reddy, A. et al. Genetic and Functional Drivers of Diffuse Large B Cell Lymphoma. Cell 171, 481–494 e415 (2017). 10.1016/j.cell.2017.09.027

51 Laude, M. C. et al. First-line treatment of double-hit and triple-hit lymphomas: Survival and tolerance data from a retrospective multicenter French study. Am J Hematol 96, 302–311 (2021). 10.1002/ajh.26068

52 Desai, S. H. et al. Improved Survival of R/R Double Hit/Triple Hit Lymphoma in the Era of CD19 Chimeric Antigen T Cell (CART) Therapy. Blood 142 (2023). 10.1182/blood-2023-183001

53 Maiti, A. et al. Outcomes of relapsed or refractory acute myeloid leukemia after frontline hypomethylating agent and venetoclax regimens. Haematologica 106, 894–898 (2021). 10.3324/haematol.2020.252569

54 Nishida, Y. et al. Stem-Cell Enriched Cellular Hierarchy of TP53 Mutant Acute Myeloid Leukemia Is Vulnerable to Targeted Protein Degradation of c-MYC. Blood 142 (2023). 10.1182/blood-2023-174938

55 Sallman, D. A. et al. TP53 mutations in myelodysplastic syndromes and secondary AML confer an immunosuppressive phenotype. Blood 136, 2812–2823 (2020). 10.1182/blood.2020006158

56 Garralda, E. et al. MYC targeting by OMO-103 in solid tumors: a phase 1 trial. Nat Med (2024). 10.1038/s41591-024-02805-1

57 Prochownik, E. V. & Wang, H. Lessons in aging from Myc knockout mouse models. Front Cell Dev Biol 11, 1244321 (2023). 10.3389/fcell.2023.1244321

58 Uy, G. L. et al. Clinical Activity of CC-90009, a Cereblon E3 Ligase Modulator and First-in-Class GSPT1 Degrader, As a Single Agent in Patients with Relapsed or Refractory Acute Myeloid Leukemia (R/R AML): First Results from a Phase I Dose-Finding Study. Blood 134 (2019). 10.1182/blood-2019-123966

59 Gavory, G. et al. Development of MRT-2359, an orally bioavailable GSPT1 molecular glue degrader, for the treatment of lung cancers with MYC-induced translational addiction. Cancer Research 83 (2023). 10.1158/1538-7445.Am2023-3449

60 Lagadinou, E. D. et al. BCL-2 inhibition targets oxidative phosphorylation and selectively eradicates quiescent human leukemia stem cells. Cell Stem Cell 12, 329–341 (2013). 10.1016/j.stem.2012.12.013

61 Simsek, T. et al. The distinct metabolic profile of hematopoietic stem cells reflects their location in a hypoxic niche. Cell Stem Cell 7, 380–390 (2010). 10.1016/j.stem.2010.07.011

62 Zeng, A. G. X. et al. A cellular hierarchy framework for understanding heterogeneity and predicting drug response in acute myeloid leukemia. Nat Med 28, 1212–1223 (2022). 10.1038/s41591-022-01819-x

63 Nishida, Y. et al. Targeting Co-Regulatory Feedback Loop of MYC and GSPT1 By MYC/GSPT1 Dual Degradation Is Synthetic Lethal in Combination with Ven/Aza in TP53 Mutant Acute Myeloid Leukemia Stem and Progenitor Cells. Blood 144, 2755 (2024). 10.1182/blood-2024-207812

64 Casey, S. C., Baylot, V. & Felsher, D. W. The MYC oncogene is a global regulator of the immune response. Blood 131, 2007–2015 (2018). 10.1182/blood-2017-11-742577

65 Casey, S. C. et al. MYC regulates the antitumor immune response through CD47 and PD-L1. Science 352, 227–231 (2016). 10.1126/science.aac9935

66 Franssen, L. E. et al. Cereblon loss and up-regulation of c-Myc are associated with lenalidomide resistance in multiple myeloma patients. Haematologica 103, e368–e371 (2018). 10.3324/haematol.2017.186601

67 Bird, S. & Pawlyn, C. IMiD resistance in multiple myeloma: current understanding of the underpinning biology and clinical impact. Blood 142, 131–140 (2023). 10.1182/blood.2023019637

## References

1 Nishida, Y. et al. Enhanced TP53 reactivation disrupts MYC transcriptional program and overcomes venetoclax resistance in acute myeloid leukemias. Sci Adv 9, eadh1436 (2023). 10.1126/sciadv.adh1436

2 Pan, R. et al. Synthetic Lethality of Combined Bcl-2 Inhibition and p53 Activation in AML: Mechanisms and Superior Antileukemic ESicacy. Cancer Cell 32, 748–760 e746 (2017). 10.1016/j.ccell.2017.11.003

3 Ishizawa, J. et al. FZR1 loss increases sensitivity to DNA damage and consequently promotes murine and human B-cell acute leukemia. Blood 129, 1958–1968 (2017). 10.1182/blood-2016-07-726216

4 Piya, S. et al. Targeting the NOTCH1-MYC-CD44 axis in leukemia-initiating cells in T-ALL. Leukemia 36, 1261–1273 (2022). 10.1038/s41375-022-01516-1

5 Shima, H., Nakayasu, M., Aonuma, S., Sugimura, T. & Nagao, M. Loss of the MYC gene amplified in human HL-60 cells after treatment with inhibitors of poly(ADP-ribose) polymerase or with dimethyl sulfoxide. Proc Natl Acad Sci U S A 86, 7442–7445 (1989). 10.1073/pnas.86.19.7442

6 Lee, W. H. et al. JAK pathway induction of c-Myc critical to IL-5 stimulation of cell proliferation and inhibition of apoptosis. J Cell Biochem 106, 929–936 (2009). 10.1002/jcb.22069

7 Wang, T., Wei, J. J., Sabatini, D. M. & Lander, E. S. Genetic screens in human cells using the CRISPR-Cas9 system. Science 343, 80–84 (2014). 10.1126/science.1246981

8 Heckl, D. et al. Generation of mouse models of myeloid malignancy with combinatorial genetic lesions using CRISPR-Cas9 genome editing. Nat Biotechnol 32, 941–946 (2014). 10.1038/nbt.2951

9 Labun, K. et al. CHOPCHOP v3: expanding the CRISPR web toolbox beyond genome editing. Nucleic Acids Res 47, W171–W174 (2019). 10.1093/nar/gkz365

10 Baran, N. et al. Inhibition of mitochondrial complex I reverses NOTCH1-driven metabolic reprogramming in T-cell acute lymphoblastic leukemia. Nat Commun 13, 2801 (2022). 10.1038/s41467-022-30396-3

11 Panina, S. B. et al. Novel mitochondria-targeting compounds selectively kill human leukemia cells. Leukemia 36, 2009–2021 (2022). 10.1038/s41375-022-01614-0

12 Tiziani, S. et al. Optimized metabolite extraction from blood serum for 1H nuclear magnetic resonance spectroscopy. Anal Biochem 377, 16–23 (2008). 10.1016/j.ab.2008.01.037

13 DiNardo, C. D. et al. Glutaminase inhibition in combination with azacytidine in myelodysplastic syndromes: a phase 1b/2 clinical trial and correlative analyses. Nat Cancer (2024). 10.1038/s43018-024-00811-3

14 Lu, X. et al. Metabolomics-based phenotypic screens for evaluation of drug synergy via direct-infusion mass spectrometry. iScience 25, 104221 (2022). 10.1016/j.isci.2022.104221

15 Lu, M. J. et al. SLC25A51 decouples the mitochondrial NAD(+)/NADH ratio to control proliferation of AML cells. Cell Metab 36, 808–821 e806 (2024). 10.1016/j.cmet.2024.01.013

16 Wang, Y., Parsons, L. R. & Su, X. AccuCor2: isotope natural abundance correction for dual-isotope tracer experiments. Laboratory Investigation 101, 1403–1410 (2021). doi:10.1038/s41374-021-00631-4

17 Ishizawa, J. et al. Mitochondrial ClpP-Mediated Proteolysis Induces Selective Cancer Cell Lethality. Cancer Cell 35, 721–737 e729 (2019). 10.1016/j.ccell.2019.03.014

18 Subramanian, A. et al. Gene set enrichment analysis: a knowledge-based approach for interpreting genome-wide expression profiles. Proc Natl Acad Sci U S A 102, 15545–15550 (2005). 10.1073/pnas.0506580102

19 Davis, M. W. & Jorgensen, E. M. ApE, A Plasmid Editor: A Freely Available DNA Manipulation and Visualization Program. Front Bioinform 2, 818619 (2022). 10.3389/fbinf.2022.818619

20 Cao, X. et al. Analysis of 3760 hematologic malignancies reveals rare transcriptomic aberrations of driver genes. Genome Med 16, 70 (2024). 10.1186/s13073-024-01331-6

21 Zou, Z., Ohta, T. & Oki, S. ChIP-Atlas 3.0: a data-mining suite to explore chromosome architecture together with large-scale regulome data. Nucleic Acids Res 52, W45–W53 (2024). 10.1093/nar/gkae358

22 Shimada, K., Bachman, J. A., Muhlich, J. L. & Mitchison, T. J. shinyDepMap, a tool to identify targetable cancer genes and their functional connections from Cancer Dependency Map data. Elife 10 (2021). 10.7554/eLife.57116

